# A novel antifolate suppresses growth of FPGS-deficient cells and overcomes methotrexate resistance

**DOI:** 10.1101/2023.02.26.530079

**Authors:** Felix van der Krift, Dick W. Zijlmans, Rhythm Shukla, Ali Javed, Panagiotis I. Koukos, Laura L.E. Schwarz, Elpetra P.M. Timmermans-Sprang, Peter E.M. Maas, Digvijay Gahtory, Maurits van den Nieuwboer, Jan A. Mol, Ger J. Strous, Alexandre M.J.J. Bonvin, Mario van der Stelt, Edwin J.A. Veldhuizen, Markus Weingarth, Michiel Vermeulen, Judith Klumperman, Madelon M. Maurice

## Abstract

Cancer cells make extensive use of the folate cycle to sustain increased anabolic metabolism. Multiple chemotherapeutic drugs interfere with the folate cycle, including methotrexate and 5-fluorouracil that are commonly applied for the treatment of leukemia and colorectal cancer (CRC), respectively. Despite high success rates, therapy-induced resistance causes relapse at later disease stages. Depletion of folylpolyglutamate synthase (FPGS), which normally promotes intracellular accumulation and activity of both natural folates and methotrexate, is linked to methotrexate and 5-fluorouracil resistance and its association with relapse illustrates the need for improved intervention strategies. In this study, we characterize a novel antifolate (C1) that, like methotrexate, potently inhibits dihydrofolate reductase (DHFR) and downstream one-carbon metabolism. Contrary to methotrexate, however, C1 displays optimal efficacy in FPGS-deficient contexts, due to decreased competition with intracellular folate concentrations for interaction with DHFR. Indeed, we show that FPGS-deficient patient-derived CRC organoids display enhanced sensitivity to C1-induced growth inhibition, while FPGS-high CRC organoids are more sensitive to methotrexate. Our results thus argue that polyglutamylation-independent antifolates can be applied to exert selective pressure on FPGS-deficient cells during chemotherapy, employing a vulnerability created by polyglutamylation deficiency.

## Introduction

Antifolates constitute a subclass of antimetabolites that have been applied as chemotherapeutic agents for decades (Stine et al., 2022; Wilson et al., 2014). In mammals, folate is an essential vitamin that functions as co-factor for the one-carbon cycle, which is among the most highly and specifically upregulated pathways in cancer (Nilsson et al., 2014). The enzyme dihydrofolate reductase (DHFR) activates folates by reducing inactive, oxidized dihydrofolate (DHF) to tetrahydrofolate (THF), which is interconverted to 5,10-methylene-THF, 10-formyl-THF and 5-methyl-THF by various enzymes of the one-carbon cycle. THF functions as a carrier for serine-and glycine-derived one-carbon units and is indispensable for supplying substrates to enzymatic pathways required for *de novo* nucleotide synthesis, NAD(P)H generation, ATP synthesis, amino acid homeostasis and tRNA modification (Ducker & Rabinowitz, 2017; Yang & Vousden, 2016; Zheng & Cantley, 2019). Folate metabolism is compartmentalized at the subcellular level, with similar reaction steps occurring in the cytosol and mitochondria, which is essential to maintain folate integrity (Zheng et al., 2018). Import of folate from the extracellular environment into the cytosol is mediated via the membrane-bound solute carriers SLC19A1 and SLC46A1 or via clathrin-mediated endocytosis of folate receptors (FOLR1-3) (Zhao et al., 2009, 2011; Zheng & Cantley, 2019). Once imported, folates are polyglutamylated by folylpolyglutamate synthase (FPGS), which catalyzes the addition of negatively charged glutamate residues to prevent efflux and promote intracellular accumulation (Lawrence et al., 2014; Osborne et al., 1993). Due to the crucial role of DHFR in maintaining sufficient concentrations of THF to drive anabolic metabolism, multiple DHFR inhibitors are clinically available for the treatment of neoplastic and autoimmune diseases, including the commonly used antifolate methotrexate. Next to targeting DHFR, the one-carbon cycle is inhibited by drugs directed at thymidylate synthase (TYMS), like 5-fluorouracil (5-FU), which is used as first-line therapy for colorectal cancer (CRC) (Stine et al., 2022).

Methotrexate is widely used to treat tumors, often in combination with other chemotherapeutics (Stine et al., 2022). Despite the high success rate of methotrexate treatment, therapy resistance presents a major problem, for example in pediatric acute lymphoblastic leukemia (ALL), where relapses occur in 20% of patients (Nguyen et al., 2008). Acquired chemoresistance observed in relapsed patients suggests that initial chemotherapy drives selection of drug-resistant clones (Li et al., 2020; Yu et al., 2020). At the cellular level, resistance to antifolates develops through various mechanisms that cause decreased cellular import, decreased retention or increased export (Fotoohi et al., 2009; Zarou et al., 2021; Zhao & Goldman, 2003). Methotrexate is an FPGS substrate and its polyglutamylation is required for efficient intracellular retention and the selective targeting of tumor cells (Fabre et al., 1984; Rots et al., 1999). Studies in cell lines and cancer cells derived from relapsed patients showed that methotrexate-induced FPGS-deficiency develops through transcriptional downregulation or the acquirement of inactivating mutations (Fotoohi et al., 2009; Li et al., 2020; Liani et al., 2003; Stark et al., 2009; Wojtuszkiewicz et al., 2016; Yu et al., 2020). Two recent studies showed that at least 8% of relapsed ALL patients obtained FPGS mutations upon methotrexate therapy (Li et al., 2020; Yu et al., 2020). Moreover, the contribution of FPGS-deficiency to methotrexate resistance may be underestimated, as over 50% of relapsed patients displays transcriptional *FPGS* downregulation (Li et al., 2020; Yu et al., 2020). Furthermore, FPGS-deficiency has been linked to resistance to 5-FU in models of CRC (Sohn et al., 2004), illustrating the need for novel intervention strategies to prevent relapses caused by drug-resistant, polyglutamylation-deficient cells.

Here, we report on the mechanistic characterization of a 2,4-diaminopyrimidine–derivative that we called compound 1 (C1), a novel and highly potent antifolate that targets FPGS-deficient cells. Functional comparison with methotrexate suggests that, although their cellular targets are similar, individual cancer cell lines display up to a 50-fold difference in sensitivity to both drugs. Cellular thermal shift assays reveal that the formation of inhibitor-DHFR complexes is strongly dependent on intracellular FPGS expression levels. We demonstrate that cells with low FPGS expression are prone to DHFR inhibition by polyglutamylation-independent antifolates, such as C1 and pyrimethamine. Using patient-derived CRC organoids that display either FPGS deficiency or overexpression, we confirm that C1 selectively inhibits growth of FPGS-deficient cells, while cells with high FPGS expression acquire sensitivity to methotrexate. Our results show that polyglutamylation-independent antifolates like C1 exert selective pressure on FPGS-deficient cells during chemotherapy and thus may be applied to prevent tumor evolution towards a methotrexate-resistant subtype. In comparison with trimetrexate, the lipophilic and FPGS-independent derivative of methotrexate, C1 has increased potency towards polyglutamylation-deficient cells. We anticipate that C1’s structure may serve as a template for development of improved non-classical antifolates.

## Results

### Identification of a 2,4-diaminopyrimidine-based compound as a novel DHFR ligand

In a screen of compounds with potential antineoplastic activity (Van der Velden et al., 2022), we identified a 2,4-diaminopyrimidine-derivative (C1, also referred to as BM001 (Van der Velden et al., 2022)) that selectively inhibited growth of a subset of cancer cell lines (Supplementary Table 1). We noted that C1 shares structural features with non-classical antifolates like pyrimethamine (Figure 1A). As C1 is structurally divergent from the classical antifolate methotrexate (Figure 1A), we compared the growth inhibitory activity of both compounds towards a panel of cancer cell lines listed in the Cancer Cell Line Encyclopedia (Ghandi et al., 2019). Remarkably, IC_50_ values between C1 and methotrexate did not correlate (Pearson r = 0.058) and a subset of cell lines even displayed up to 50-fold differences in sensitivity (Figure 1B). These results thus suggest that, despite its predicted antifolate activity, C1 inhibits tumor cell growth by a mode of action different from methotrexate.

**Figure 1.**
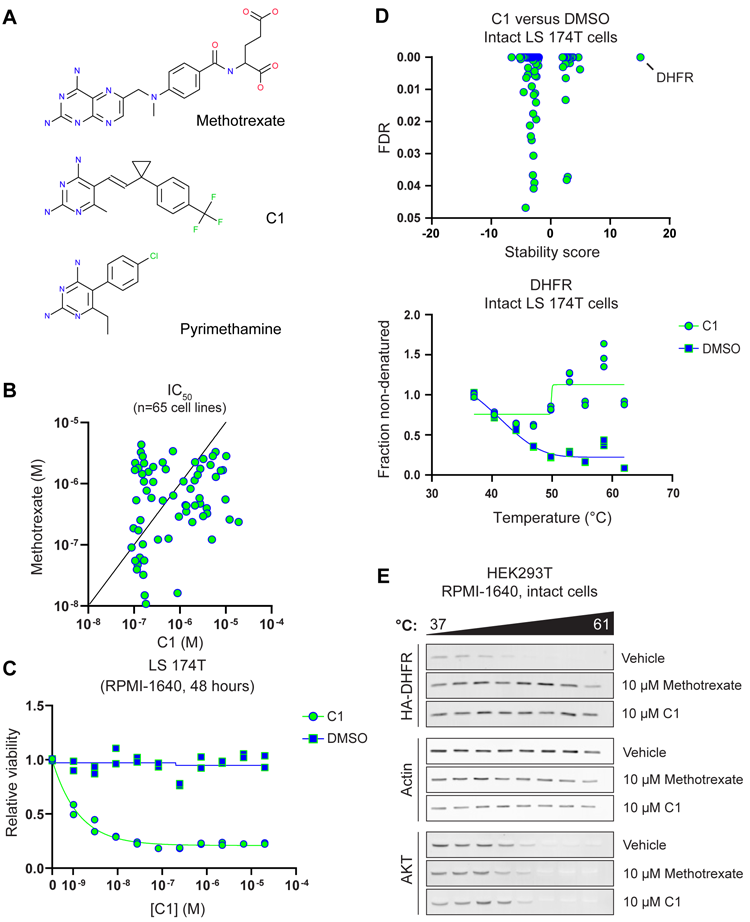
Identification of compound C1 as a novel DHFR ligand. **(A)** Structures of Methotrexate, Pyrimethamine and C1, a novel 2,4-diaminopyrimidine antifolate. **(B)** Comparison of C1 and Methotrexate IC_50_ values for a panel of cancer cell lines. Methotrexate IC_50_ values were retrieved from the Genomics of Drug Sensitivity in Cancer database and determined using a Syto60 viability assay after 72h treatment. C1 IC_50_ values were determined using a Sulforhodamine B viability assay after 72h treatment. **(C)** CellTiter-Glo viability assay on C1-or vehicle-treated LS 174T cells after 48 hours treatment. Data were collected for n=2 biological replicates. **(D)** Top panel: thermal proteome profile of intact LS 174T cells treated with 10 μM C1 for 1 hour, including stability score and false discovery rate (FDR) of all hits and candidates. Negative stability scores represent proteins that are destabilized by C1 treatment, positive stability scores represent proteins that are stabilized by C1 treatment. Bottom panel: melting curve for DHFR in 10 μM C1-or DMSO-treated intact LS 174T cells. **(E)** Western-blot analysis of a representative cellular thermal shift assay on intact HEK293T cells overexpressing HA-DHFR treated with 10 μM Methotrexate or C1 for 1 hour.

To understand the underlying growth inhibitory mechanism, we aimed to identify key cellular proteins targeted by C1. Based on the presence of a nitrogenous base, we first hypothesized that C1 may act as an ATP-competitive kinase inhibitor. However, we did not detect *in vitro* kinase inhibition using 100 nM C1 in a kinome-wide screen (Supplementary Figure 1). Next, we interrogated C1-protein interactions by applying mass spectrometry-based thermal proteome profiling (TPP) (Mateus et al., 2020; Molina et al., 2013; Savitski et al., 2014). We employed the LS 174T CRC cell line that is sensitive to C1 treatment (Figure 1C). LS 174T cells were either mock-treated or treated for 1 hour with 10 μM C1, after which cells were subjected to TPP. A remarkable C1-mediated thermal shift was observed for DHFR, indicating that C1 primarily acts as a DHFR ligand, as suspected based on its 2,4-diaminopyrimidine group (Figure 1D). Additionally, we observed C1-mediated stabilization for TYMS (Supplementary Table 2), a known nM affinity target of polyglutamylated methotrexate (Huber et al., 2015). This finding was unexpected, as C1 lacks a glutamate group for FPGS-mediated polyglutamylation. We therefore speculate that TYMS may instead be stabilized by the accumulation of dUMP, due to 5,10-methylene-THF depletion that occurs in DHFR-inhibited cells (Brown et al., 2023; Ducker & Rabinowitz, 2017; Yang & Vousden, 2016; Zheng & Cantley, 2019).

To validate C1 as a novel DHFR ligand, we overexpressed human DHFR (hDHFR) bearing an N-terminal HA-tag in HEK293T cells and performed a cellular thermal shift assay (CETSA) after treatment for 1 hour with either 10 μM C1 or methotrexate (Molina et al., 2013). Western-blot analysis revealed that both methotrexate and C1 stabilize HA-DHFR when compared to DMSO-treated cells, while the thermal stability of control proteins (AKT and Actin) remained unaffected by both drug treatments (Figure 1E). These findings thus confirm that treatment with C1, like methotrexate, stabilizes DHFR in live cells, suggesting a direct interaction. Based on our combined findings, we conclude that C1 is a novel hDHFR ligand, similar to methotrexate. The observed differential growth inhibitory effects of C1 and methotrexate on various cancer cell lines, however, indicates that their mode of action does not fully overlap or may be context-dependent.

### C1 inhibits DHFR and downstream events in folate-mediated one-carbon metabolism

To analyze pharmacological inhibition of DHFR, we measured *in vitro* activity of recombinant DHFR in the presence of various concentrations C1 or methotrexate, using excess concentrations of DHF and NADPH substrate. Methotrexate inhibited DHFR activity close to 100% at all tested concentrations, in line with the previously reported 6 pM K_i_ of methotrexate towards mouse DHFR at 100 nM DHF (Figure 2A) (Piper et al., 1985). C1 also inhibited *in vitro* DHFR activity, although with a higher IC_50_ of approximately 1 μM in the presence of 50 μM DHF, suggesting that C1 binds to hDHFR with a lower affinity than methotrexate (Figure 2A). Next, we applied isothermal titration calorimetry (ITC) to determine the dissociation constants (K_d_) of DHFR to C1 and methotrexate (Table 1 and Supplementary Figure 2A). On average, the K_d_ of DHFR for methotrexate was 5 nM, while the K_d_ of DHFR for C1 was 28 nM, indicating that the affinity of C1 for DHFR is weaker than methotrexate. Addition of 125 µM NADPH did not affect binding of both C1 and methotrexate (Supplementary Figure 2B). These results show that, like methotrexate, C1 directly inhibits DHFR, albeit with lower affinity.

**Figure 2.**
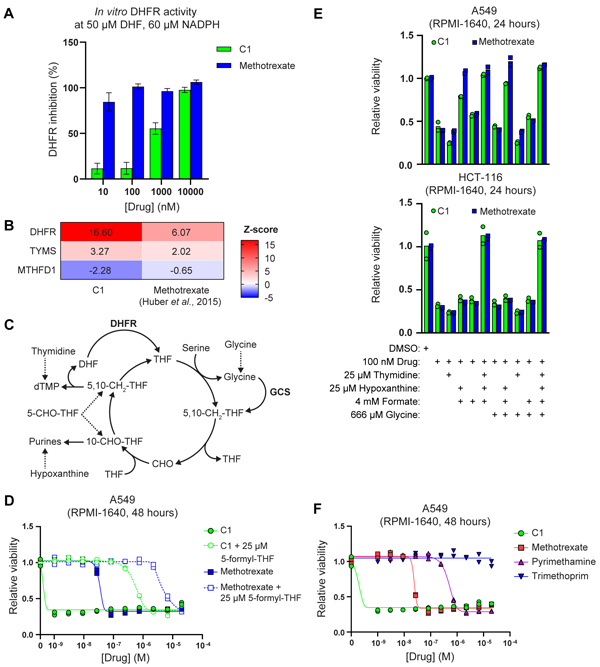
C1 and methotrexate inhibit DHFR and folate-mediated one-carbon metabolism. **(A)** *In vitro* inhibition of DHFR activity by C1 and Methotrexate, at 50 μM dihydrofolate (DHF) and 60 μM NADPH. Data are shown as mean ± s.d. and were collected in n=2 technical replicates. Results are representative of 3 independent experiments. **(B)** Thermal proteome profiling analysis of one-carbon metabolism-related proteins in cells treated with C1 or Methotrexate, including Z-score transformed stability scores. Positive Z-scores represent stabilized proteins, negative Z-scores represent destabilized proteins. The thermal proteome profile for C1 was determined at 10 μM in LS 174T cells (Figure 1D). The thermal proteome profile for methotrexate was retrieved from Huber et al. (2015) and determined at 10 μM in K562 cells. **(C)** Simplified schematic representation of tetrahydrofolate (THF)-mediated one-carbon transfer reactions involved in purine and thymidine synthesis. Rescue interventions are indicated with dashed lines. Abbreviations: 5,10-methylenetetrahydrofolate (5,10-CH2-THF), formate (CHO), 10-formyl-tetrahydrofolate (10-CHO-THF), 5-formyl-tetrahydrofolate (5-CHO-THF), glycine cleavage system (GCS). **(D)** CellTiter-Glo viability analysis of C1-or methotrexate-treated A5 cells after 48 hours treatment, in the absence or presence of 25 μM 5-formyl-tetrahydrofolate. Data were collected in n=2 biological replicates. **(E)** CellTiter-Glo viability analysis of C1-or methotrexate-treated A549 or HCT-116 cells after 24h treatment, including rescue treatments with one-carbon cycle metabolites. Data were collected in n=2 biological replicates. **(F)** CellTiter-Glo viability analysis of C1-, methotrexate-, pyrimethamine-, or trimethoprim-treated A549 cells after 48h treatment. Data were collected in n=2 biological replicates.

**Table 1.** Dissociation constants (K_d_) of DHFR to C1 and methotrexate in absence of NADPH, determined by isothermal titration calorimetry.

| Experiment | Average $K_d$ (± s.d.) (nM) | Measured $K_d$ range (nM) | Replicates |
| --- | --- | --- | --- |
| DHFR to C1 | 28.10 ± 18.38 | 15.1 – 41.1 | 2 |
| DHFR to methotrexate | 4.75 ± 0.80 | 4.18 – 5.32 | 2 |

Next to monitoring drug-protein interactions, thermal proteome profiling may be used to assess drug-induced alterations in metabolite-protein interactions within cells (Huber et al., 2015). We hypothesized that inhibition of DHFR would indirectly lead to thermal shifts of enzymes catalyzing THF-dependent reactions as a result of altered substrate availability. To obtain insight into the metabolic consequences of C1 treatment, we used PANTHER statistical overrepresentation analysis to identify biological processes and pathways linked to alterations in proteome stability of C1-treated LS 174T cells (Mi et al., 2019). Proteins involved in various anabolic processes were significantly overrepresented in the thermal proteome profile of C1-treated cells, including pathways linked to nucleotide biosynthesis, transcription and translation (Table 2). We also observed C1-induced destabilization of C-1-tetrahydrofolate synthase (MTHFD1) (Figure 2B), which suggests loss of interactions, indicative of decreased substrate availability and thus corroborates C1-mediated inhibition of cytosolic 5,10-methylene-THF and 10-formyl-THF production. To compare the TPP of C1-treated LS 174T cells with methotrexate, we analyzed the published TPP of K562 cells treated with 10 μM methotrexate for 3 hours and performed z-score transformation on both datasets to enable comparison (Huber et al., 2015). Both C1 and methotrexate significantly stabilized DHFR and TYMS, while C1, but not methotrexate, did cause significant destabilization of MTHFD1 (Figure 2B).

**Table 2.** Thermal proteome profiling reveals various anabolic processes and pathways affected by C1 treatment in LS 174T cells. PANTHER GO-slim Biological Process and PANTHER Pathways statistical overrepresentation test on (de)stabilized hits and candidates listed in Figure 1D. The table includes fold enrichment and false discovery rate (FDR).

| PANTHER GO-Slim Biological Process | Fold enrichment | FDR |
| --- | --- | --- |
| positive regulation of RNA polymerase II transcription preinitiation complex assembly (GO:0045899) | > 100 | 5.71E-08 |
| pyrimidine nucleotide biosynthetic process (GO:0006221) | 52.81 | 9.47E-04 |
| proteasome assembly (GO:0043248) | 45.26 | 2.03E-02 |
| ribosomal large subunit assembly (GO:0000027) | 41.33 | 9.34E-07 |
| RNA polymerase II preinitiation complex assembly (GO:0051123) | 34.44 | 2.54E-05 |
| nucleoside monophosphate metabolic process (GO:0009123) | 25.01 | 6.53E-03 |
| ribosomal small subunit biogenesis (GO:0042274) | 17.6 | 6.96E-05 |
| ribonucleoside triphosphate biosynthetic process (GO:0009201) | 12.43 | 8.08E-03 |
| cytoplasmic translation (GO:0002181) | 11.74 | 9.48E-03 |
| purine ribonucleotide biosynthetic process (GO:0009152) | 10.93 | 7.38E-04 |
| DNA-dependent DNA replication (GO:0006261) | 10.56 | 1.36E-02 |
| translational elongation (GO:0006414) | 8.73 | 5.67E-07 |
| proteasome-mediated ubiquitin-dependent protein catabolic process (GO:0043161) | 6.53 | 1.13E-03 |
| oxoacid metabolic process (GO:0043436) | 3.99 | 1.08E-02 |
| PANTHER Pathways | Fold enrichment | FDR |
| Sulfate assimilation (P02778) | > 100 | 7.34E-03 |
| Tetrahydrofolate biosynthesis (P02742) | 63.37 | 1.62E-02 |
| ATP synthesis (P02721) | 52.81 | 1.93E-02 |
| Formyltetrahydroformate biosynthesis (P02743) | 45.26 | 2.25E-02 |
| Cell cycle (P00013) | 43.21 | 4.40E-07 |
| Ubiquitin proteasome pathway (P00060) | 31.17 | 1.59E-12 |
| DNA replication (P00017) | 15.33 | 2.21E-02 |

To compare the cellular effects of C1 and methotrexate in more detail, we treated A549 cells with a titration of both drugs for 48 hours, including a rescue with 25 μM 5-formyl-THF, a metabolite that can be converted to 5,10-methylene-THF and 10-formyl-THF, independently of DHFR function (Figure 2C) (Ducker & Rabinowitz, 2017). Using an ATP-based cell viability assay, A549 cells displayed high sensitivity to C1 with a sub-nM IC_50_, while the IC_50_ for methotrexate was around 50 nM. The IC_50_ of both drugs increased approximately 400-fold upon 5-formyl-THF supplementation (Figure 2D), suggesting that both drugs impair one-carbon transfer via the folate cycle, presumably through inhibition of DHFR. To assess whether C1 and methotrexate inhibit specific reactions of the one-carbon cycle, we treated A549 and HCT-116 cells with 100 nM drug and investigated rescue effects by supplementation with the one-carbon cycle metabolites thymidine, hypoxanthine, formate and glycine (Figure 2C). C1-and methotrexate-treated A549 and HCT-116 cells were rescued by a combination of thymidine and hypoxanthine (Figure 2E; Supplementary Figure 3A), suggesting that both drug treatments deplete intracellular 5,10-methylene-THF and 10-formyl-THF pools, required for purine and thymidine synthesis (Figure 2C) (Ducker & Rabinowitz, 2017; Lawrence et al., 2014; Yang & Vousden, 2016; Zheng et al., 2018). Supplementation with formate or glycine, which can be used by MTHFD1 to generate cytosolic 10-formyl-THF or by the glycine cleavage system to generate mitochondrial 5,10-methylene-THF, respectively, did not rescue viability of C1-and methotrexate-treated cells (Figure 2E). These findings suggest that depletion of intracellular THF limits these reactions (Figure 2C) (Ducker & Rabinowitz, 2017; Yang & Vousden, 2016). The nutrient sensing branch of the mechanistic target of rapamycin complex 1 (mTORC1) signaling network responds to intracellular purine levels, leading to mTORC1 inhibition upon antifolate treatment (Emmanuel et al., 2017; Hoxhaj et al., 2017). To compare the effects of C1 and methotrexate on purine sensing by mTORC1, we treated A549 cells overnight with both drugs, followed by a 2.5 hours supplementation of adenosine and guanosine to restore intracellular purine levels. Western-blot analysis of the mTORC1 substrate 4E-BP1 revealed mTORC1 inhibition by C1 and methotrexate, which could be partially restored by adenosine but not guanosine supplementation (Supplementary Figure 3B). This observation is in accordance with the report that mTORC1 is inhibited by short-term depletion of adenylates and suggests that methotrexate and C1 have similar effects on intracellular purines and associated mTORC1 activity (Hoxhaj et al., 2017).

Notwithstanding these *in vitro* results, we observed that a subset of cancer cell lines (including A549) display over 50-fold higher sensitivity to C1 than methotrexate, pyrimethamine and trimethoprim (Figure 2F). These observations suggest that inhibition of DHFR activity *in vitro* does not necessarily reflect the intracellular situation within live cells, where folate concentrations are expected to be lower (Bailey et al., 2015). Combined, our results show that C1 is a novel DHFR inhibitor that interferes with THF-mediated one-carbon transfer reactions required for *de novo* nucleotide synthesis, DNA replication, transcription and translation. Furthermore, our results show that C1 is less potent in inhibiting hDHFR than methotrexate, while some cell lines display a 50-fold higher sensitivity to C1, emphasizing the pivotal role of intracellular context on evaluating cellular drug responses.

### C1 and methotrexate inhibit DHFR by a partially overlapping binding mode

Our *in vitro* results might suggest that C1 and methotrexate may have different DHFR binding modes. To compare DHFR binding by C1 with methotrexate and natural folates, we generated docking models of the hDHFR-NADPH-C1 complex by using published crystal structures of hDHFR in complex with 2,4-diaminopyrimidine compounds and we compared our models with structures of hDHFR in complex with natural folates or methotrexate (Table 3). The hDHFR active site cleft transitions from an open (Figure 3A) to a closed conformation upon binding to ligand (Figure 3B-C) or to methotrexate (Figure 3D) (Bhabha et al., 2013; Cody et al., 2005). We observed a similar closed conformation in the docking model of the hDHFR-NADPH-C1 complex, although both open (Figure 3E) and closed (Figure 3F) conformations of Phe31 were predicted energetically favorable. hDHFR is known to bind folate and 5,10-dideaza-THF through formation of hydrogen bonds with Glu30, Asn64 and Arg70 as well as hydrophobic interactions with Phe31, which causes the active site to adopt a closed conformation (Figure 3G-H) (Bhabha et al., 2013). These contacts are conserved in the hDHFR-NADPH-methotrexate complex. In addition, methotrexate forms a hydrogen bond with Tyr121 (Figure 3I-J). Our predicted hDHFR-NADPH-C1 complex indicates that hydrogen bonds with Tyr121 and Glu30, as well as hydrophobic contacts with Phe31, are conserved (Figure 3K). By contrast, Asn64 and Arg70 do not participate in complex formation of DHFR with C1 (Figure 3L), providing a potential explanation for the higher IC_50_ value of C1 compared to methotrexate (Figure 3H, 3J; Figure 2A). Taken together, our results suggest that the binding mode of C1 to hDHFR partially overlaps with methotrexate, while displaying an overall reduced binding surface. These results explain the lower affinity of C1 compared to methotrexate observed in our *in vitro* experiments. Of note, our simulation system was not biased with any specific restraints that could recreate the hydrogen bond network that is observed with folates and methotrexate, indicating a similar binding mode for C1.

**Table 3.**
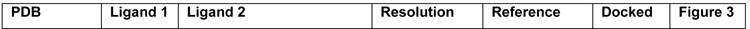
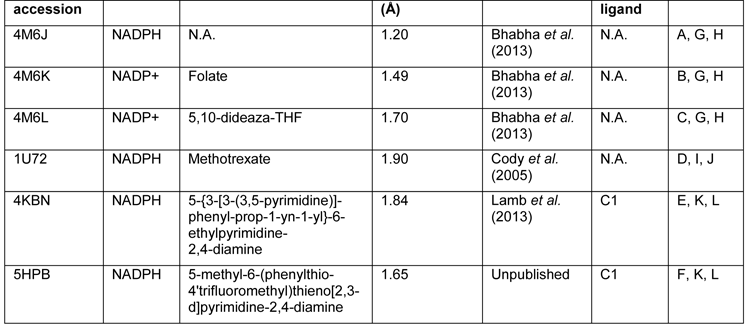
Crystal structures used for generation of docking models of the DHFR-NADPH-C1 complex and comparison with methotrexate and natural folates.

| PDB | Ligand 1 | Ligand 2 | Resolution | Reference | Docked | Figure 3 |
| --- | --- | --- | --- | --- | --- | --- |

| accession |  |  | (A) |  | ligand |  |
| --- | --- | --- | --- | --- | --- | --- |
| 4M6J | NADPH | N.A. | 1.20 | Bhabha *et al.* (2013) | N.A. | A, G, H |
| 4M6K | NADP+ | Folate | 1.49 | Bhabha *et al.* (2013) | N.A. | B, G, H |
| 4M6L | NADP+ | 5,10-dideaza-THF | 1.70 | Bhabha *et al.* (2013) | N.A. | C, G, H |
| 1U72 | NADPH | Methotrexate | 1.90 | Cody *et al.* (2005) | N.A. | D, I, J |
| 4KBN | NADPH | 5-{3-[3-(3,5-pyrimidine)]-phenyl-prop-1-yn-1-yl}-6-ethylpyrimidine-2,4-diamine | 1.84 | Lamb *et al.* (2013) | C1 | E, K, L |
| 5HPB | NADPH | 5-methyl-6-(phenylthio-4'trifluoromethyl)thieno[2,3-d]pyrimidine-2,4-diamine | 1.65 | Unpublished | C1 | F, K, L |

**Figure 3.**
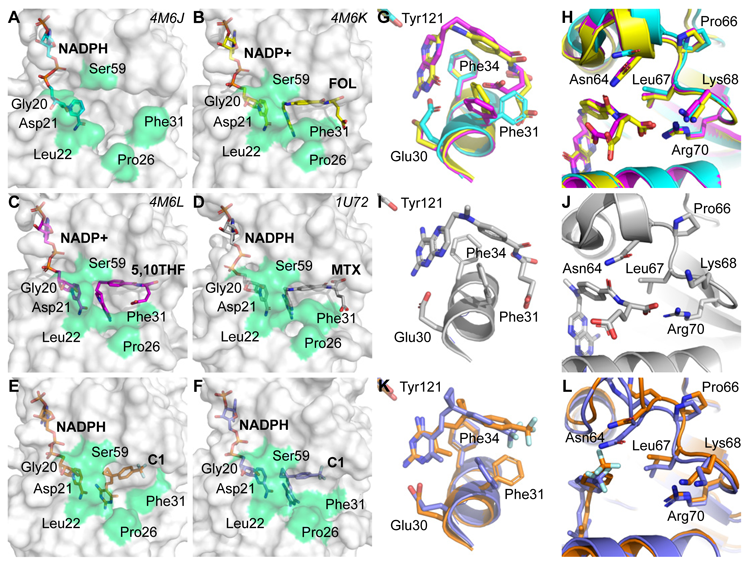
DHFR binding modes of C1 and methotrexate are predicted to be partially overlapping. Surface representation of **(A)** crystal structure of hDHFR in complex with NADPH, **(B)** crystal structure of hDHFR in complex with NADP+ and folic acid (FOL), **(C)** crystal structure of hDHFR in complex NADP+ and 5,10-dideaza-THF (5,10THF), **(D)** crystal structure of hDHFR in complex with NADPH and methotrexate (MTX), (E) docking pose of C1 on hDHFR (PDB accession 4M6J) and **(F)** docking pose of C1 on hDHFR (PDB accession 5HPB). Residues involved in opening and closing of the active site are highlighted in green and were adopted from Bhabha et al. (2013). Active site crystal structures of the **(G-H)** hDHFR-NADPH (cyan), hDHFR-NADPH-folic acid (yellow), hDHFR-NADP+-5,10-dideaza-THF (magenta) and **(I-J)** hDHFR-NADPH-methotrexate complexes. **(K-L)** Docking poses of the hDHFR-NADPH-C1 complexes (orange is PDB accession 4KBN, purple is PDB accession 5HPB).

### FPGS expression determines intracellular DHFR engagement by antifolates

Our results reveal that C1 has a higher affinity for DHFR than DHF, but a lower affinity than methotrexate. Nevertheless, a subset of cell lines displays up to 50-fold higher sensitivity to C1, emphasizing that cellular context is a major determinant of antifolate sensitivity. A key regulatory element that is missing in *in vitro* assays is the presence of membrane-encapsulated subcellular compartments. Methotrexate and natural folates do not diffuse across membranes due to their charged glutamate residue(s), but are actively imported by the FOLR pathway, SLC19A1 and SLC46A1, while intracellular accumulation is promoted by FPGS-mediated polyglutamylation (Zheng & Cantley, 2019). Conversely, polyglutamates can be hydrolyzed to monoglutamates by the lysosomal or secreted enzyme γ-glutamyl-hydrolase (GGH), which is thought to facilitate membrane transport of methotrexate and natural folates (Zheng & Cantley, 2019). These properties categorize Methotrexate as a classical antifolate (Gangjee et al., 2007, 2008). By contrast, hydrophobic compounds such as C1 and pyrimethamine cross membranes by passive diffusion and do not depend on FPGS activity for cellular retention, classifying them as non-classical antifolates.

In line with suggestions in literature, we hypothesized that the FPGS-dependent concentrations of intracellular folate determine sensitivity of cells to classical or non-classical antifolates like C1 (Fabre et al., 1984; Fotoohi et al., 2009; Li et al., 2020; Liani et al., 2003; Rots et al., 1999; Stark et al., 2009; Wojtuszkiewicz et al., 2016; Yu et al., 2020; Zarou et al., 2021; Zhao & Goldman, 2003). To address this issue, we employed data from the Cancer Cell Line Encyclopedia (Ghandi et al., 2019), to calculate correlation coefficients between polyglutamylation-associated gene expression and sensitivity (IC_50_) to C1 or methotrexate, for a panel of cell lines (Supplementary Figure 4A). In addition, we employed two larger datasets to investigate how polyglutamylation-associated gene expression correlates with cellular sensitivity for pyrimethamine and methotrexate (Supplementary Figure 4B). Notably, *FPGS* expression positively correlated with sensitivity to methotrexate (Supplementary Figure 4B), while displaying an inverse correlation with C1 sensitivity (Supplementary Figure 4A). This analysis did not reveal a relationship between sensitivity to C1 and expression of *GGH* (Supplementary Figure 4A), while *GGH* expression correlated with resistance to methotrexate (Supplementary Figure 4B). These findings thus further corroborate FPGS-deficiency as a mechanism of methotrexate resistance and suggest that FPGS determines whether cells display sensitivity to C1 or methotrexate. Interestingly, this methotrexate therapy-induced escape route of cancer cells thus may uncover a cellular vulnerability to C1.

To analyze the effect of FPGS on intracellular DHFR binding by antifolates, we applied CETSA in a semi-quantitative experimental setup for measuring intracellular DHFR-antifolate binding. We used intact HEK293T cells overexpressing DHFR with and without FPGS and tested a drug dilution series, while keeping a fixed, denaturing temperature (Figure 4; Supplementary Figure 5). C1 potently stabilized DHFR at 10 nM and higher concentrations, in line with the 28 nM K_d_ of the C1-DHFR complex determined by ITC (Figure 4A, C; Table 1). Methotrexate and pyrimethamine also stabilized DHFR, although at higher concentrations than C1, indicative of decreased intracellular complex formation with DHFR (Figure 4A, C). In contrast with the 5 nM K_d_ of the methotrexate-DHFR complex determined by ITC (Table 1), 10 nM of methotrexate does not lead to intracellular DHFR stabilization (Figure 4A, C), supporting the view that the presence of membrane-encapsulated intracellular compartments limits the action of this hydrophilic molecule. Overexpression of FPGS decreased intracellular DHFR stabilization by C1 and pyrimethamine, which indicates decreased intracellular complex formation and confirms that FPGS overexpression impairs DHFR binding by non-classical antifolates (Figure 4B-C). By contrast, FPGS overexpression mediated a shift in DHFR stabilization to lower concentrations of methotrexate (100 nM) (Figure 4B-C). Combined, these results suggest that FPGS overexpression increases the intracellular concentration of natural folates, which likely competes with DHFR binding of polyglutamylation-independent antifolates (C1 and pyrimethamine). On the other hand, methotrexate accumulates more efficiently in FPGS-high cells, shifting optimal DHFR binding to lower methotrexate concentrations. We conclude that inhibitory activity of C1 inversely correlates with intracellular FPGS levels, suggesting that C1 may be applied to selectively target FPGS-deficient cancer cells.

**Figure 4.**
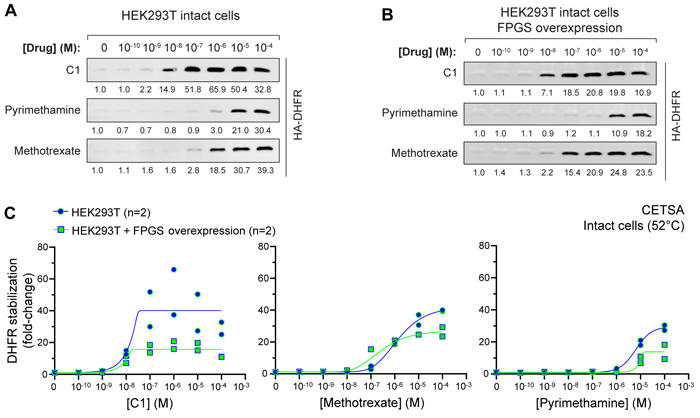
FPGS expression determines the degree of intracellular DHFR binding by C1, methotrexate and pyrimethamine. Representative Western-blot analyses of the cellular thermal shift assays on intact HEK293T cells **(A)** overexpressing HA-DHFR or **(B)** HA-DHFR and FPGS-FLAG. Cells were incubated with a drug concentration range for 90 minutes followed by a cellular thermal shift assay at 52.0 °C. Relative HA-DHFR intensity, normalized for vehicle control (0M drug), is indicated below each panel. **(C)** Quantification and comparison of relative HA-DHFR stabilization in intact HEK293T cells with and without FPGS-FLAG overexpression.

### C1 selectively suppresses growth of FPGS-deficient cells

Next to mediating methotrexate resistance, loss of FPGS is a known mechanism of resistance towards 5-FU treatment in CRC cells (Sohn et al., 2004), which may uncover a vulnerability for C1 treatment. To address this issue, we employed patient-derived CRC organoids characterized by FPGS deficiency to analyze the relationship between polyglutamylation and therapeutical efficiency of antifolates and 5-FU (Van de Wetering et al., 2015). RT-qPCR analysis of patient 6-derived tumor organoids (P6T) confirms that this line displays transcriptional downregulation of *FPGS* expression compared to healthy colon organoids (HCO). At the same time, the expression of *GGH*, which catalyzes hydrolysis of folate polyglutamates, is increased in P6T organoids (Supplementary Figure 6). These findings indicate that P6T organoids have a decreased capacity to polyglutamylate FPGS-dependent antifolates, which is expected to decrease the intracellular concentration and efficacy of these drugs. Indeed, a viability assay of P6T organoids at 7 days after start of treatment revealed that this organoid line is approximately 3-fold more sensitive to C1 than to methotrexate, in line with our model that FPGS-deficiency creates a vulnerability to C1 (Supplementary Figure 7).

To further test how efficacy of different antifolate classes depends on cellular polyglutamylation capacity, we used transposase-mediated integration of doxycycline-inducible FPGS, GGH and empty vector (EV) overexpression constructs, thereby including independent expression cassettes for fluorescent mNeongreen, mCherry or NLS-TagBFP reporters, respectively. Immunofluorescence analysis confirmed that FPGS and GGH were overexpressed when organoids were cultured in the presence of doxycycline (Supplementary Figure 8). P6T organoids overexpressing FPGS, GGH and EV were cultured in the presence of 100 and 500 nM C1 or 200 and 500 nM methotrexate and outgrowth was analyzed on day 8 by imaging fluorescent organoids, followed by a viability assay. The lower methotrexate concentration was set at 200 nM to compensate for the decreased sensitivity of P6T organoids for methotrexate compared to C1 (Supplementary Figure 7). P6T organoids overexpressing EV showed a dose-dependent decrease in outgrowth upon treatment with either C1 or methotrexate (Figure 5A), which was even more apparent in the viability assay (Figure 5B). An increased sensitivity of the chemical viability assay may be explained by the fact that alterations in intracellular metabolite concentrations (fast) may occur before outgrowth (slow) is impaired. Organoids overexpressing FPGS showed a comparable dose-dependent decrease in outgrowth during C1 treatment as observed for non-FPGS-overexpressing cells, but were strongly sensitized to methotrexate treatment, which decreased organoid outgrowth by 80% (Figure 5A). Viability analysis further revealed that FPGS overexpression in organoids mediated a rescue from treatment with 100 nM C1 (p < 0.0001), but not 500 nM C1 (Figure 5B). These findings are in line with our model that FPGS prevents DHFR-C1 complex formation by enhancing the intracellular concentrations of folate, which can be outcompeted by increasing doses of C1. P6T organoids overexpressing GGH showed a dose-dependent decrease in outgrowth upon both drug treatments (Figure 5A). In contrast to FPGS, overexpression of GGH created a minor rescue from treatment with 100 nM C1 (p = 0.0348), but not 500 nM C1 (Figure 5B). For 200 nM methotrexate, GGH overexpression caused a more significant rescue (p < 0.0001), which was not observed for 500 nM methotrexate (Figure 5B). Combined, these results reveal that FPGS-deficient P6T organoids display sensitivity to C1 while being more resistant to methotrexate, which is reverted upon re-introduction of FPGS, indicating that FPGS-deficiency sensitizes to FPGS-independent antifolates like C1. The role of GGH appears to be more complex since it decreases toxicity of both methotrexate and C1, however, and may involve differential effects on the import of natural folates and drug compounds.

**Figure 5.**
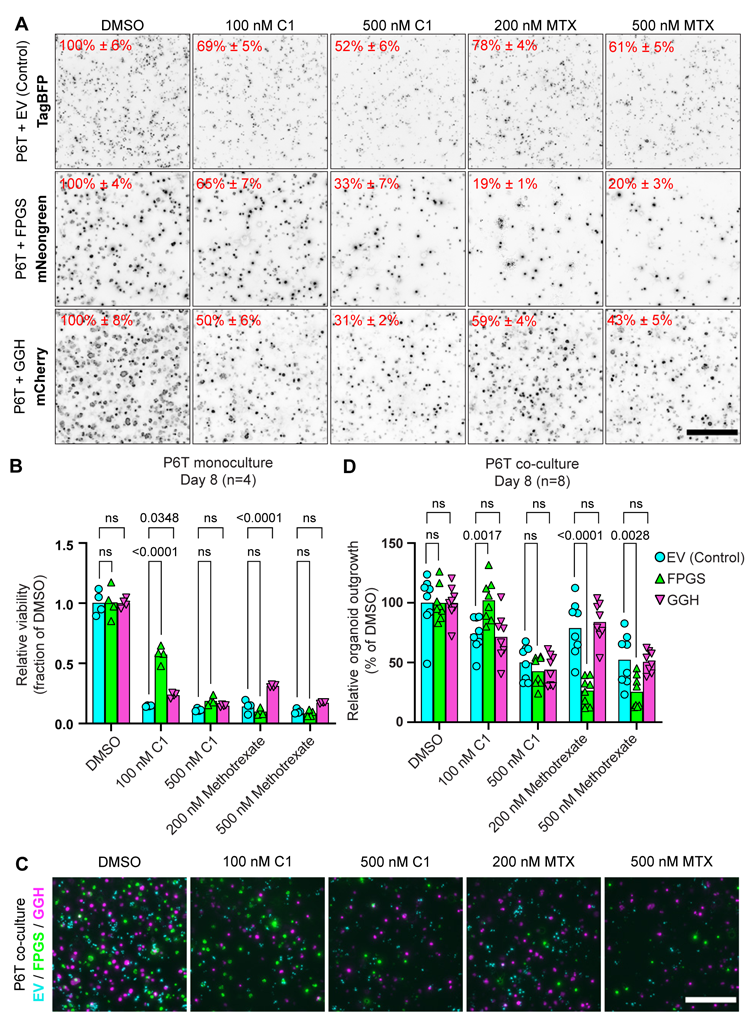
FPGS-deficiency causes methotrexate resistance, but creates a vulnerability to C1. **(A)** Representative widefield fluorescence images of P6T organoids overexpressing doxycycline-inducible constructs with a fluorescent reporter after 8 days of drug treatment. Overexpression of empty vector IRES-TagBFP (EV), FPGS-FLAG IRES-mNeongreen (FPGS) or GGH-FLAG IRES-mCherry (GGH) was combined with various drug treatments for 8 days. Organoids were treated with DMSO (vehicle control), 100 or 500 nM C1 and 200 or 500 nM Methotrexate (MTX). Relative organoid outgrowth is measured as % surface area of DMSO ± 95% confidence interval. Images are Z-stack projections of deconvoluted widefield images. Scalebar represents 500 μm. Data were collected in n=2 independent experiments with biological duplicates. **(B)** CellTiter-Glo viability analysis of P6T organoids overexpressing empty vector, FPGS or GGH after 8 days of drug treatment. Data were collected in n=2 independent experiments with biological duplicates. Viability of FPGS-and GGH-overexpressing organoids was compared with empty vector-overexpressing organoids using two-way ANOVA with Dunnett’s multiple comparisons test. **(C)** Representative widefield fluorescence images of co-cultured P6T organoids overexpressing EV (blue), FPGS (green) or GGH (red) after 8 days of drug treatment. Overexpression was combined with various drug treatments for 8 days. Images are Z-stack projections of deconvoluted widefield images. Scalebar represents 500 μm. Data was collected in n=2 independent experiments with biological quadruplicates. **(D)** Quantification of fluorescence surface area in widefield images of co-cultured P6T organoids overexpressing EV (blue), FPGS (green) or GGH (red) after 8 days of drug treatment. Analysis was performed on Z-stack projections of deconvoluted widefield images. Relative surface area of FPGS-and GGH-overexpressing organoids was compared with empty vector-overexpressing organoids using two-way ANOVA with Dunnett’s multiple comparisons test. Data was collected in n=2 independent experiments with biological quadruplicates.

Next, we examined if clinically approved polyglutamylation-independent antifolates also preferentially target FPGS-deficient cells, like C1. To address this point, we performed viability assays using FPGS-deficient P6T organoids and patient 26-derived CRC organoids (P26T) that display similar FPGS expression levels as healthy colon organoids (Van de Wetering et al., 2015). We assessed the response to C1, methotrexate, the polyglutamylation-independent methotrexate-derivative trimetrexate and the TYMS inhibitor 5-FU. Compared to P26T organoids, P6T organoids were relatively insensitive to methotrexate, trimetrexate and 5-FU, but displayed sensitivity to C1. Overall, P6T organoids were 3-fold more sensitive to C1 than to methotrexate and trimetrexate (Supplementary Figure 7). This suggests that C1 classifies as a more potent polyglutamylation-independent antifolate than trimetrexate for the treatment of FPGS-deficient cells, although we cannot rule out that trimetrexate activity and/or resistance may occur through alternative mechanisms.

### C1 overcomes methotrexate resistance of FPGS-deficient tumor organoids

In ALL, primary tumors were reported to contain persister clones that survive initial chemotherapy treatment and grow out at later timepoints to cause therapy resistance and relapse (Li et al., 2020; Yu et al., 2020). In methotrexate-treated patients, FPGS-deficiency may develop through inactivating mutations or transcriptional downregulation resulting from copy number alterations or promoter deletions (Li et al., 2020; Yu et al., 2020). We hypothesized that C1 treatment may prevent outgrowth of FPGS-deficient clones in a heterogeneous tumor and thereby overcome methotrexate resistance. To mimic tumor heterogeneity, we co-cultured FPGS-deficient P6T organoid lines overexpressing EV (control), FPGS or GGH and evaluated their relative outgrowth during treatment with various concentrations of C1 or methotrexate. Outgrowth was measured on day 8 by imaging fluorescent reporters and quantification of organoid surface area (Figure 5C-D). While 100 nM C1 suppressed growth of FPGS-deficient organoids (overexpressing EV or GGH), FPGS overexpression caused full resistance to 100 nM C1 in co-culture conditions (p = 0.0017) (Figure 5C-D). FPGS-mediated resistance was lost at 500 nM C1, in line with our previous results (Figure 5A-B). Conversely, 200 and 500 nM methotrexate suppressed growth of FPGS-overexpressing organoids (p < 0.0001 and p = 0.0028, respectively), while FPGS-deficient organoids were partially resistant (Figure 5C-D). P6T organoids overexpressing GGH responded identically to all tested antifolate treatments as organoids overexpressing EV, which suggests that the relationship between antifolate efficacy and polyglutamylation is primarily shaped by FPGS, not GGH. Taken together, our results show that FPGS deficiency creates a vulnerability for C1, while cells with functional FPGS are sensitive to methotrexate. Our findings indicate that a regimen in which methotrexate-based chemotherapy is alternated with C1 treatment will suppress outgrowth of FPGS-deficient, methotrexate-resistant persister clones which may cause disease relapse at later stages.

## Discussion

Methotrexate has been successfully used for chemotherapy, but its therapeutic index is limited by off-target toxicity, including bone marrow suppression, damage to gastrointestinal epithelia and hepatotoxicity (Takami et al., 1995; Wilson et al., 2014). These problems fostered efforts to discover new antifolates with improved therapeutic index, illustrated by 7 clinically approved and 15 DHFR inhibitors currently undergoing clinical or preclinical evaluation (Cuthbertson et al., 2021). In ALL, methotrexate treatment has a high initial success rate, but therapy-induced selection of drug-resistant cells decreases its therapeutic index, which involves depletion of FPGS activity in at least 8% of pediatric ALL patients (Li et al., 2020; Nguyen et al., 2008). Accordingly, polyglutamylation-independent antifolates were acknowledged as potential agents to overcome methotrexate resistance (Cho et al., 2007; Kim et al., 2013; Sohn et al., 2004). Here, we report on the characterization of C1, a novel polyglutamylation-independent antifolate. C1 has been studied as a pesticide (patent WO9820878A1), but recently regained interest for use in human subjects after showing potent inhibition of tumor growth and limited toxicity in a mouse model, where it was studied for potential effects on growth hormone signaling (Van der Velden et al., 2022). To our knowledge, our study is the first in-depth characterization of C1 as antifolate in human tissue-derived model systems, although a group of compounds with similarities to C1 has been studied *in vitro* using purified hDHFR (Algul et al., 2011). We find that C1, like methotrexate, selectively binds and inhibits DHFR, impairs one-carbon transfer to anabolic reactions and inhibits cellular growth. In contrast to methotrexate, C1 displays strongest DHFR inhibition in FPGS-deficient cellular contexts and outperforms the FPGS-independent antifolate trimetrexate in polyglutamylation-deficient tumor organoids.

To interrogate the cellular interaction partners of C1, we employed thermal proteome profiling (TPP) (Mateus et al., 2020; Molina et al., 2013; Savitski et al., 2014). The results unambiguously identify DHFR as the primary target of C1. Furthermore, TPP of C1-treated cells revealed destabilization of multiple one-carbon cycle enzymes, presumably through decreased substrate availability as a result of THF depletion. These observations are in accordance with C1 acting as an antimetabolite and indicate that the folate pathway is the main cellular process affected by C1 intervention. Additionally, our findings support the use of thermal proteome profiling for identification of protein engagement by cellular metabolites (Huber et al., 2015). By using a modified CETSA protocol similar to two-dimensional thermal proteome profiling (Becher et al., 2016), we performed semi-quantitative measurements of intracellular DHFR-antifolate complexes, which allowed us to demonstrate the effect of cellular FPGS levels on DHFR binding to both classical and non-classical antifolates. Overexpression of FPGS decreased intracellular DHFR binding by C1 and pyrimethamine, but not methotrexate. Importantly, C1 concentrations needed for optimal intracellular DHFR stabilization were in the range of 10-100 nM, matching the 28 nM K_d_ that we determined for the DHFR-C1 complex by ITC.

Loss of FPGS expression links to treatment resistance against 5-FU (Sohn et al., 2004), the most commonly applied chemotherapy in CRC. 5-FU efficacy typically depends on ternary complex formation with TYMS and 5,10-methylene-THF (Longley et al., 2003). 5-FU-mediated complex formation is compromised upon FPGS depletion due to a decrease in intracellular THF levels (Longley et al., 2003; Sohn et al., 2004), thus offering a clinically relevant condition to investigate whether polyglutamylation-deficient cancer cells display an acquired sensitivity to C1. To obtain proof-of-concept, we employed patient-derived CRC organoids characterized FPGS-deficiency or FPGS-overexpression. Our results confirm that FPGS-deficient CRC organoids are relatively insensitive to 5-FU, methotrexate and trimetrexate, but acquire a vulnerability for treatment with C1, in line with our model.

Methotrexate-resistant persister clones are cells that have an intrinsic or acquired insensitivity to therapy and are present at the start or formed during initial chemotherapy treatment and cause relapse in later stages (Li et al., 2020). By co-culturing different organoids, we mimic the heterogeneous situation that occurs during initial chemotherapy treatment of primary tumors in patients. Our results show that methotrexate treatment confers a competitive advantage to FPGS-deficient cells, while C1 treatment favors outgrowth of FPGS-overexpression cells. This finding has two key implications. First, polyglutamylation deficiency creates a vulnerability with sufficient therapeutic window for exploitation by C1. Second, FPGS-dependent and polyglutamylation-independent antifolates are not interchangeable and therapy based on a single agent will inevitably favor the outgrowth of persister clones. Our work therefore warrants a combination or alternating therapy of FPGS-dependent and polyglutamylation-independent antifolates to suppress growth of both the FPGS-deficient and FPGS-competent populations. Viability studies show that C1 has improved capabilities to inhibit expansion of FPGS-deficient CRC organoids compared to trimetrexate, a lipophilic and FPGS-independent derivative of methotrexate. It is unclear why C1 outperforms trimetrexate and additional biochemical studies are needed to compare relevant properties of C1 with other non-classical antifolates to address this issue, including affinity for hDHFR and subcellular distribution. C1 was found serendipitously and we anticipate that its structure may serve as a template for the development of improved non-classical antifolates that have optimal properties for selective inhibition of FPGS-deficient, methotrexate resistant cells. Apart from FPGS-deficiency, methotrexate resistance develops through decreased import and increased export, for example by inactivating mutations in SLC19A1 or overexpression of ABCG2 (Zarou et al., 2021; Zhao & Goldman, 2003). Considering the lipophilic nature of C1, it would be interesting to test if lipophilic antifolates like C1 may also selectively suppress growth of cells that are methotrexate-resistant as a result of altered membrane transport.

In conclusion, we identified a highly potent polyglutamylation-independent antifolate that selectively suppresses growth of methotrexate-or 5-FU-resistant, FPGS-deficient tumor cells. Our results show that FPGS-deficiency causes an exploitable vulnerability to C1 and warrant a combination therapy of FPGS-dependent and-independent antifolates to prevent expansion of persister cells and overcome methotrexate resistance.

## Materials & Methods

### Cell culture

HEK293T, HCT-116, LS 174T and A549 were cultured in RPMI-1640 (Sigma-Aldrich) supplemented with 10% fetal bovine serum (Bodinco), 2 mM UltraGlutamine (Lonza), 100 units/ml penicillin and 100 µg/ml streptomycin (Sigma-Aldrich). Cells were cultured at 37°C in 5% CO_2_ and regularly checked for mycoplasma.

### Organoid culture

P6T and P26T human colorectal cancer organoids were established in a previous study (Van De Wetering et al., 2015) and obtained following a material transfer agreement with Hubrecht Organoid Technology (Utrecht, The Netherlands). Organoids were cultured in advanced DMEM/F12 medium (ThermoFisher Scientific), supplemented with penicillin-streptomycin (Sigma-Aldrich), 10 mM HEPES (ThermoFisher Scientific), 1X Glutamax (ThermoFisher Scientific), B27 (ThermoFisher Scientific), 10 mM nicotinamide (Sigma-Aldrich), 1.25 mM N-acetylcysteine (Sigma-Aldrich), 10% v/v Noggin-conditioned medium, 50 ng/ml human EGF (Peprotech), 500 nM A83-01 TGF-ß type 1 receptor inhibitor (Tocris) and 10 µM SB202190 P38 MAPK inhibitor (Sigma-Aldrich). Organoids were maintained in Cultrex BME (R&D systems) and dissected using TrypLE (ThermoFisher) during passaging. Medium was supplemented with 10 µM Y-27632 ROCK inhibitor (Selleck Chemicals) after splitting.

### Organoid electroporation

Electroporation of organoids was performed as previously described using a NEPA21 electroporator (Fujii et al., 2015). 7.2 µg PiggyBac-CMV overexpression construct, 7.2 µg rTTA-IRES-Hygro and 5.2 µg PiggyBac transposase were used to generate organoid lines with inducible overexpression of FPGS, GGH and empty vector. 100 µg/ml Hygromycin B was used for selection of electroporated organoids.

### Organoid viability assays

For viability assays upon treatment with antifolates, advanced DMEM/F12 medium was replaced with MEM (ThermoFisher Scientific) supplemented with 1X MEM non-essential amino acids (ThermoFisher Scientific) and 5 µg/L vitamin B12 (Sigma-Aldrich). 1 µg/ml doxycycline was added to induce overexpression of FPGS, GGH and empty vector constructs. Before seeding, organoids were dissociated using TrypLE (ThermoFisher Scientific) and passed through a 100 µm cell strainer. 1000X drug dilutions were added 2 days after seeding. For monoculture viability assays, 12500 cells/well were seeded in a 96-well plate containing 10 µl BME. CellTiter-Glo luminescent cell viability assay (Promega) was mixed 1:1 with culture medium and 200 µl/well was used to quantify viability of monocultures. For co-culture viability assays, 12500 cells/well of each line were seeded in a 96-well plate containing 50 µl BME matrix topped off with 20 µl empty BME. Outgrowth of organoids in co-culture was analyzed using a Leica THUNDER widefield microscope equipped with a 10X HC PL FLUOTAR objective (NA=0.32) and large volume computational clearing deconvolution algorithm. Organoid surface area was quantified from Z-stack maximum projections using FIJI MorphoLibJ Morphological Segmentation plugin combined with particle analysis. Brightfield images were obtained on an EVOS M5000 imaging system (ThermoFisher Scientific).

### Plasmids and antibodies

Human HA-DHFR was subcloned from human cDNA to pcDNA4/TO by PCR using Q5 High Fidelity 2X mastermix (NEB). Human FPGS-FLAG and GGH-FLAG were subcloned from cDNA clones MHS6278-202755815 (Horizon Discovery) and MHS6278-202757326 (Horizon Discovery), respectively, to pcDNA4/TO and PiggyBac-CMV-MCS-IRES-mCherry and PiggyBac-CMV-MCS-IRES-mNeongreen by PCR using Q5 High Fidelity 2X mastermix (NEB). PiggyBac-CMV-MCS-IRES-NLS-TagBFP was used as empty vector control. All constructs were sequence verified. The following primary antibodies were used for Western blotting (WB) and immunofluorescence (IF): rat anti-HA (Roche, 11867423001), mouse anti-FLAG (Sigma-Aldrich, F3165), rabbit anti-4E-BP1 (Cell Signaling, 9452), mouse anti-TOM20 (BD transduction laboratories, 612278), mouse anti-LAMP1 (BD Pharmingen, 555798), rabbit anti-Akt (Cell Signaling, 9272) and mouse anti-Actin (MP Biomedicals, 691001). Primary antibodies were diluted according to manufacturer’s instructions. Secondary antibodies for WB and IF were diluted 1:5000 and 1:300, respectively, and obtained from Rockland, LI-COR or Invitrogen.

### Immunofluorescence and confocal microscopy of organoids

Organoids were released from extracellular matrix using dispase (Sigma-Aldrich) at 37°C for 30 minutes. Organoids were fixed in 0.1M phosphate buffer containing 4% paraformaldehyde for 1 hour. Washing and antibody stainings were perfomed in 4-well µ-Slides (Ibidi), using PBS containing 0.2% Triton X-100, 1% DMSO and 1% bovine serum albumin. 1 µg/ml DAPI was used to stain nuclei. Organoids were mounted using mounting medium (Ibidi) and analyzed using a Zeiss LSM700 confocal microscope equipped with 63X plan-apochromat oil immersion objective (NA=1.40).

### Cell viability assays

5000 cells/well were seeded in a 96-well plate and drugs diluted in full culture medium were added the next day. Viability was quantified with CellTiter-Glo luminescent cell viability assay (Promega) according to manufacturer’s instructions. Luciferase activity was measured on a Berthold Centro LB960 luminometer. Drug sensitivity of a panel of cancer cell lines for C1 was determined at OncoLead (Karsfeld, Germany), using a Sulforhodamine B viability assay after 72 hours of treatment (Figure 1B; Supplementary Table 1; Supplementary Figure 4).

### Drugs and rescue metabolites

Pyrimethamine (Sigma-Aldrich), methotrexate (Selleck Chemicals), trimethoprim (Sigma-Aldrich), trimetrexate hydrochloride (CI-898; Santa Cruz) were dissolved in DMSO. Rescue agents were dissolved in water unless indicated otherwise and treatments with hypoxanthine (dissolved in 67% formate; Sigma-Aldrich), folinic acid (5-formyl-THF; Sigma-Aldrich), thymidine (Sigma-Aldrich), glycine (Sigma-Aldrich), adenosine (Sigma-Aldrich) and guanosine (Sigma-Aldrich) were performed as indicated. Compound C1 was synthesized by Specs Compound Handling (Zoetermeer, The Netherlands) as described in patent WO2021078995A1.

### Reverse-transcriptase quantitative PCR

Reverse-transcriptase quantitative PCR was performed as previously described (Omerzu et al., 2019). RNA was isolated using a Qiagen RNeasy Mini Kit (Qiagen). Turbo DNAse (ThermoFisher Scientific) was used to remove genomic DNA. cDNA was synthesized using a iScript cDNA synthesis kit (Bio-Rad). Human *FPGS* and *GGH* primers were previously published (Driehuis et al., 2020). Other primers used: CTTTTGCGTCGCCAG (*GAPDH* forward), TTGATGGCAACAATATCCAC (*GAPDH* reverse). *FPGS* and *GGH* expression was calculated relative to *GAPDH* using the 2^-ΔCt^ method.

### Cellular thermal shift assays

HEK293T cells were cultured in RPMI-1640 to 50% confluency in 100 mm plates and subsequently transfected with 6 µg/dish pcDNA4/TO_HA-DHFR or 5 µg/dish pcDNA4/TO_HA-DHFR + 5 µg/dish pcDNA4/TO_FPGS-FLAG. Transfections were performed using polyethylenimine (PEI) and medium was refreshed after 4 hours incubation with the transfection mix. Cellular thermal shift assay (CETSA) was performed as described before (Zheng et al., 2018) and samples were heated to 37, 39, 42.3, 46.4, 51.9, 56.1, 59 and 61°C in an S1000 thermal cycler (Bio-Rad). For CETSA with dose-response, drug incubations were performed in PCR tubes, using 100 µl cell suspension per condition. 100X drug stocks were diluted to 1X and incubated for 90 minutes. CETSA results were analyzed by mixing the cell extracts with sample buffer and subsequent Western blotting.

### Cell lysis and Western blotting

Cells were washed once with ice-cold PBS and lysed in RIPA buffer (25 mM Tris pH 7.6, 150 mM NaCl, 1% IGEPAL CA-630 (NP-40 alternative), 1% sodium deoxycholate, 0.1% SDS, 50 mM NaF, 1 mM PMSF, 10 μg/ml Leupeptin, 10 μg/ml Aprotinin). Lysates were centrifuged at 15000 × g for 15 minutes at 4°C. Soluble fraction was isolated and mixed with 5X sample buffer (350 mM Tris pH 6.8, 10% SDS, 20% glycerol, 2.5% 2-mercaptoethanol, 0.025% bromophenol blue) to a final concentration of 1X. Samples were denatured at 95°C for 10 minutes. Western blotting was performed under standard procedures using SDS-PAGE to resolve samples, followed by transfer to Immobilon-FL PVDF membrane (Millipore). Membranes were blocked with Odyssey blocking buffer (LI-COR) diluted 1:1 in TBS. Primary and secondary antibodies were diluted in TBS + 0.05% Tween20. Membranes were imaged on an Amersham Typhoon NIR laser scanner (GE Healthcare).

### Thermal proteome profiling

Thermal proteome profiling was done as previously described (Becher et al., 2018). In brief, cells were harvested after 1 hour of treatment with 10 μM compound C1 or DMSO, washed with PBS and 10 aliquots, each of 1×10^6^ cells in 100 μL PBS, were distributed in a 96-well PCR plate. After centrifugation (300 × g for 3 minutes) and removal of most of the supernatant (80 μL), each aliquot was heated for three minutes to a different temperature (37°C, 40.4°C, 44°C, 46.9°C, 49.8°C, 52.9°C, 55.5°C, 68.6°C, 62°C, 66.3°C) in a PCR machine (Agilent SureCycler 8800) followed by 3 minutes at room temperature. Cells were lysed with 30 μL ice-cold lysis buffer (final concentration 0.8% NP-40, 1.5mM MgCl2, protease inhibitors, phosphatase inhibitors, 0.4 U/μL benzonase) on a shaker (500 rpm) at 4°C for one hour. The PCR plate was then centrifuged at 300 × g for 3 minutes at 4°C to remove cell debris, and the supernatant was filtered at 300 × g for 3 minutes at 4°C through a 0.45-μm 96-well filter plate (Millipore, MSHVN4550) to remove protein aggregates. Of the flow-through, 25 μL was mixed with 2× sample buffer (180 mM Tris pH 6.8, 4% SDS, 20% glycerol, 0.1 g bromophenol blue) and kept at −20°C until prepared for mass spectrometry analysis, while the remainder was used in a BCA (ThermoFisher Scientific), to determine the protein concentration. Samples were diluted to 1 μg/μL in 1x sample buffer based on the protein concentrations in the lowest two temperatures (37°C, 40.4°C).

### MS sample preparation and measurement

Proteins were digested as previously described (Mateus et al., 2020). Briefly, 10 μg of protein (based on the protein concentrations in the lowest two temperatures) was added to a bead suspension (10 μg of beads (Thermo Fischer Scientific—Sera-Mag Speed Beads, 4515-2105-050250, 6515-2105-050250) in 10 μl 15% formic acid and 30 μl ethanol) and incubated on a shaker (500 rpm) for 15 minutes at RT. Beads were washed four times with 70% ethanol and proteins were digested overnight in 40 μl digest solution (5 mM chloroacetamide, 1.25 mM TCEP, 200 ng trypsin, and 200 ng LysC in 100 mM HEPES pH 8). Peptides were then eluted from the beads, vacuum-dried, reconstituted in 10 μl of water, and labeled for 1 hour at RT with 18 μg of TMT10plex (ThermoFisher Scientific) dissolved in 4 μl of acetonitrile (the label used for each experiment can be found in Supplementary Table 3). The reaction was quenched with 4 μl of 5% hydroxylamine, and samples were combined by temperature. Samples were acidified and desalted using StageTips (Rappsilber et al., 2007) and eluted with 2x 30 μl of buffer B (80% acetonitrile, 0.01% TFA). Samples were fractionated using the Pierce™ High pH Reversed-Phase Peptide Fractionation Kit (ThermoFisher Scientific) into 3 fractions (Fraction No. 4, 7 and 8). The flowthrough, wash and TMT wash fractions were pooled together with fraction 4. Peptides were applied to reverse-phase chromatography using a nanoLC-Easy1000 coupled online to a Thermo Orbitrap Q-Exactive HF-X. Using a 120 minutes gradient of buffer B, peptides were eluted and subjected to tandem mass spectrometry. The mass spectrometer was operated in Top20 mode and dynamic exclusion was applied for 30 seconds.

### MS data analysis

MS data were analyzed using Proteome Discoverer (ThermoFisher Scientific, version 2.2). Data were searched against the human UniProt database. Search parameters: trypsin, missed cleavages 3, peptide tolerance 10ppm, 0.02Da for MS/MS tolerance. Fixed modifications were carbamidomethyl on cysteines and TMT10plex on lysine. Variable modifications included acetylation on protein N terminus, oxidation of methionine and TMT10plex on peptide N-termini.

### Abundance and stability score calculation

The Proteome Discoverer output files were loaded into R, merged, filtered for duplicates and proteins with less than 2 unique peptides and saved in an ExpressionSet R-object. Potential batch effects were removed using limma (Ritchie et al., 2015) and data were normalized using variance stabilization, vsn strategy (Huber et al., 2002). Normalization was done for each temperature independently, to account for the decreasing signal intensity at the higher temperatures. The abundance score of each protein was calculated as the average log2 fold change at the two lowest temperatures (37°C, 40.4°C). The stability score of each protein was calculated by subtracting the abundance score from the log2 fold changes of all temperatures and calculating the sum of the resulting values. To assess the significance of abundance and thermal stability scores, we used a limma analysis, followed by an FDR analysis using the fdrtool package.

### PANTHER statistical overrepresentation analysis

From the thermal proteome profile of C1-treated LS 174T cells, all proteins displaying significant thermal shifts were used as input for PANTHER (version 16.0, released 2020-12-01) statistical overrepresentation test (released 20210224) as previously described (Mi et al., 2019). Binomial test with false discovery rate correction was used. PANTHER GO-Slim Biological Process and PANTHER Pathways were used as annotation datasets. Hierarchical clustering was applied to the PANTHER GO-Slim annotated dataset to filter out the most specific subclasses.

### Kinome-wide screening for kinase inhibition

*In vitro* screening for kinase inhibition by compound C1 was performed by Thermo Fisher Scientific (Madison, WI, United States). For the LanthaScreen kinase activity assay, fluorescein-labeled substrate was incubated with a kinase of interest and ATP to allow kinase-dependent phosphorylation. A terbium-labeled antibody was subsequently used to detect substrate phosphorylation, resulting in Förster resonance energy transfer (FRET). Time-resolved (TR) FRET was interpreted as a readout of kinase activity. For the Adapta kinase activity assay, kinase activity was reconstituted *in vitro*, followed by ADP detection using an europium-labeled antibody and an AlexaFluor647-labeled ADP-tracer, resulting in FRET in the absence of ADP production. Kinase activity produces unlabeled ADP, resulting in displacement of the labeled ADP-tracer and TR-FRET inhibition. Increased TR-FRET was interpreted as a readout of kinase inhibition. For the Z’LYTE assay, kinase activity was reconstituted *in vitro* using fluorescein-and coumarin-labeled FRET peptides as substrates. A protease cleaving non-phosphorylated peptides was subsequently added to the reaction. Proteolytic cleavage of the substrate peptides disrupts FRET and FRET inhibition was interpreted as a readout of kinase inhibition. Results were combined and visualized using Coral (Metz et al., 2018).

### In vitro DHFR activity assay

*In vitro* DHFR activity was analyzed by using a colorimetric assay kit according to manufacturer’s instructions (Sigma-Aldrich) to quantify DHFR-dependent NADPH turnover by monitoring absorbance at 340 nm. Briefly, C1 and methotrexate were dissolved to 10 mM in 30% acetonitrile + 0.1% formic acid. Drug stocks were diluted in assay buffer to final concentrations of 10-10000 nM. Absorbance at 340 nm was measured using a SmartSpec spectrophotometer (Bio-Rad) every 60 seconds for 5 minutes and quantified using kinetics software (Bio-Rad).

### Statistical analysis, correlation analysis and curve fitting

Statistical tests were performed as indicated in figure legends. All statistical analyses were performed in GraphPad Prism 9. D’Agostino & Pearson’s test was used to test for normality and lognormality. In case of small sample sizes, descriptive statistics were used to compare standard deviations for statistical tests that assume equal standard deviations. Binding and viability curves were fitted in GraphPad Prism 9, using a nonlinear regression asymmetric sigmoidal (5 parameter) model where X is concentration. Correlation between gene expression and drug sensitivity was calculated using Spearman correlation coefficient. Gene expression levels and drug sensitivity data for methotrexate and pyrimethamine were retrieved from the Genomics of Drug Sensitivity in Cancer database and the Cancer Cell Line Encyclopedia (Ghandi et al., 2019).

### Isothermal Titration Calorimetry

ITC measurements were performed in a Low Volume NanoITC (TA Instruments-Waters LLC, New Castle, DE, USA). DHFR (4.3 μM) (R&D Systems, 8456-DR-100) was prepared in assay buffer (50 mM MES hydrate, 25 mM Tris, 100 mM Nacl, 25 mM Ethanolamine, 2 mM DTT). C1 (1 µM), Methotrexate (1 µM) and β-NADPH (125 µM) were also prepared in the same buffer. 300 μL of the C1 or Methotrexate in presence or absence of β-NADPH was present in the cell (maximum cell volume is 169 μL) and 50 μL of DHFR was loaded in the syringe as titrant. At 37 °C, 2 μl of DHFR was injected into the cell every 300 seconds, except the first injection which is of 0.96 μl. All experiments were performed at 37 °C while stirring at 300 rpm. The data was analyzed with the NanoAnalyze Software (TA instruments, Asse, Belgium) and background titration of DHFR to buffer is subtracted from all thermograms.

### Generation of docking models

The protocol we followed for this system is based on our, recently published, protein-small molecule shape-restrained docking protocol (Koukos et al., 2021). For in-depth details regarding the pre-processing and docking protocol see the “Materials and Methods” section of the original publication. The main steps are summarized in the sections below.

### Template identification

First, we searched the Protein Data Bank (PDB) (Berman et al., 2000), using the SMILES string of the target compound and the FASTA sequence of the target receptor as inputs, and additionally filtering for receptors with at least one co-crystallised compound (Weininger, 1988; Weininger et al., 1989). After removing unsuitable templates (low-resolution structures, receptors only crystallised with crystallisation buffers, etc.), we extracted the template SMILES strings. Using the extracted template SMILES strings and the target compound C1 SMILES string as inputs, we compared the chemical similarity of the target compound C1 to all template compounds. For this similarity comparison we computed the Tversky similarity (biased with a weight of 0.8 for the target compound vs one of 0.2 for the template compounds) over the maximum common substructure (MCS) as identified with the rdFMCS implementation of RDKit (version 2020.09.3) (Tversky, 1977). After calculating all similarity values, we ranked the compounds by their Tversky similarity and selected the one with the highest value (closer to 1 as opposed to 0). For this modelling effort, we opted to perform the docking using the template identified via the above procedure (PDB ID: 5HPB) as well as the second-best template (PDB ID: 4KBN).

### Conformer generation

Since no 3D structures of the C1 compound were available, we opted to generate 3D conformers of the compound starting from its isomeric SMILES string: CC1=C(C(=NC(=N1)N)N)/C=C/C2(CC2) C3=CC=C(C=C3)C(F)(F)F. In total, 64 conformers were generated with RDKit using the 2020 parameter set while making use of energy minimisation and the ETKDG algorithm (Riniker & Landrum, 2015; Wang et al., 2020). We provide the ensemble of generated conformers to HADDOCK without any additional filtering.

### System preparation

We removed all crystallographic waters and all crystallisation artefacts from both template receptor structures prior to docking, while maintaining the relevant NADPH cofactor. Topologies and parameters for the cofactor and compound C1 were generated with PRODRG (version 070118.0614) (Schüttelkopf & Van Aalten, 2004).

### Generation of shape-based restraints

All docking simulations were carried out with the command-line version of HADDOCK 2.4 (January 2021 release), our integrative modelling platform, using in-house computational resources (Dominguez et al., 2003; van Zundert et al., 2016). HADDOCK integrates experimentally (or otherwise) obtained data in the docking to guide the simulation toward generating poses that satisfy the provided data. For this modelling effort we made use of restraints extracted from the shape of the template compounds. Specifically, after identifying the template receptors via the procedure laid out above, we transformed the heavy atoms of the template compounds into dummy beads and then defined ambiguous distance restraints with an upper limit of 1 Å between the shape beads and the non-hydrogen atoms of the compound to be docked. These restraints are always defined from the smaller to the larger body, so for the docking based on the 5HPB template the restraints were defined from the shape beads to any non-hydrogen atom of the compound, while for the one based on 4KBN they were defined from the non-hydrogen compound atoms to any shape bead. In addition to these shape-based restraints we also defined restraints between the non-hydrogen atoms of the NADPH cofactor and its surrounding residues meant to maintain the original geometry of the cofactor relative to its surroundings.

### Docking

For the docking, we generated 1280 models during the rigid-body stage (20 * *N_generated conformers_*) out of which the top 200 proceeded to the flexible refinement stage. We set the total number of components active during docking to 3 (the template receptor, the generated conformers and the template-based shape), disabled the systematic sampling of 180-symmetrical poses during the rigid-body stage, disabled the random removal of restraints, fixed the position of the template receptor and shape to their original positions and disabled the deletion of non-polar hydrogens. We used constant dielectric for both stages and set its value for the refinement stage to 10. We lowered the scaling of intermolecular interactions during rigid-body minimisation to 0.001 of its original value to allow the generated conformers to more easily penetrate into the binding pocket and also ignored the contribution of the vdW term during the scoring of the poses for the rigid body stage. We clustered the generated models using an RMSD cut-off value of 1.5 Å.

### Scoring

The scoring functions used for both stages are:

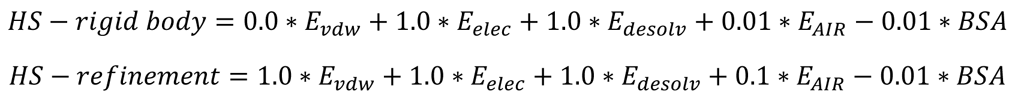

Where HS stands for HADDOCK score, Evdw, Eelec and Edesolv stand for van der Waals, Coulomb electrostatics and desolvation energies, respectively and BSA for the buried surface area. The non-bonded components of the score (Evdw, Eelec) are calculated with the OPLS forcefield (Jorgensen & Tirado-Rives, 1988). The desolvation energy is a solvent accessible surface area-dependent empirical term which estimates the energetic gain or penalty of burying specific sidechains upon complex formation (Fernández-Recio et al., 2004).

## Data availability

The mass spectrometry proteomics datasets have been deposited in the ProteomeXchange Consortium via the PRIDE partner repository (https://www.ebi.ac.uk/pride/) (Perez-Riverol et al., 2019) under the dataset identifier PXD040155.

## Acknowledgements

We thank our colleagues of the M.M.M. laboratory and Center for Molecular Medicine for fruitful discussions, feedback and suggestions; Corlinda ten Brink for assistance during imaging experiments using Cell Microscopy Core equipment; Joep Sprangers and Remco Sleiderink for providing the PiggyBac-CMV-MCS-IRES-mNeongreen plasmid; Dennis Piet, Sirik Deerenberg and Tom Speksnijder for the synthesis of C1; Nanda Sprenkels for QC, formulation and solubility studies on C1; Ingrid Jordens for providing healthy colon organoids (HCO) RNA; Michael Hadders and Joep Sprangers for critical reading of the manuscript; the members of the NWO-TTW user committee for their critical input and discussions. This work is part of the Oncode Institute, which is partly financed by the Dutch Cancer Society (KWF). This work was supported by the Netherlands Organization for Scientific Research (NWO) domain TTW, grant 16083 (to J.K., G.J.S. and J.A.M.), Zon-MW VICI grant 91815604 (to M.M.M.), Zon-MW TOP grant 91218050 (to M.M.M.) and Gravitation project IMAGINE! (to M.M.M.).

## Author contributions

F.K., D.W.Z., R.S., P.I.K., E.P.M.T.S., D.G., M.N., J.A.M., G.J.S., A.M.J.J.B., M.S., M.V., J.K. and M.M.M. conceived and designed the experiments. F.K., D.W.Z., R.S., A.J., P.I.K., L.L.E.S., E.P.M.T.S., D.G. and M.N. performed the experiments. F.K., D.W.Z., R.S., A.J., P.I.K., E.P.M.T.S., D.G., M.N., E.J.A.V., J.K. and M.M.M. analyzed the data. P.E.M.M. provided essential reagents. F.K., D.W.Z., P.I.K., R.S. and M.M.M. wrote the manuscript, which was reviewed by all authors.

## Tables and supplementary tables

**Supplementary Table 1.** C1 inhibits growth of cancer cell lines with variable potency, related to Figure 1. IC_50_ values for C1 on a panel of cancer cell lines of varying origin. IC_50_ values were determined using Sulforhodamine B viability assay after 72 hours treatment. Z-scores are calculated over all tested cell lines, using log-transformed IC_50_ values.

| Compound | Cell line | Origin | IC<sub>50</sub> (M) | Z-score |
| --- | --- | --- | --- | --- |
| C1 | NCIH82 | lung | 8.82E-08 | -1.285 |
| C1 | RDES | bone | 1.19E-07 | -1.2803 |
| C1 | UMUC3 | bladder | 1.04E-07 | -1.1919 |
| C1 | HCT116 | colon | 9.76E-08 | -1.1499 |
| C1 | SKMEL5 | skin | 1.41E-07 | -1.1259 |
| C1 | A375 | skin | 1.07E-07 | -1.1085 |
| C1 | SKHEP1 | liver | 1.12E-07 | -1.104 |
| C1 | MHHES1 | bone | 1.24E-07 | -1.0747 |
| C1 | HL-60 | hematological | 1.15E-07 | -1.0688 |
| C1 | NCIH460 | lung | 1.06E-07 | -1.0676 |
| C1 | A2780 | ovary | 1.20E-07 | -1.033 |
| C1 | MT3 | breast | 1.43E-07 | -1.0062 |
| C1 | HCT15 | colon | 1.33E-07 | -0.9624 |
| C1 | 786O | kidney | 1.37E-07 | -0.9524 |
| C1 | MDAMB468 | breast | 4.91E-07 | -0.9506 |
| C1 | A549 | lung | 1.44E-07 | -0.9497 |
| C1 | MV4-11 | hematological | 1.81E-07 | -0.938 |
| C1 | THP-1 | hematological | 1.67E-07 | -0.9192 |
| C1 | T24 | bladder | 1.54E-07 | -0.918 |
| C1 | SKLMS1 | uterus | 1.64E-07 | -0.9093 |
| C1 | MIAPACA2 | pancreas | 1.65E-07 | -0.8883 |
| C1 | C33A | endometrial | 2.06E-07 | -0.8848 |
| C1 | L-363 | hematological | 1.71E-07 | -0.8753 |
| C1 | JAR | placenta | 1.55E-07 | -0.86 |
| C1 | HT1080 | connective tissue | 1.52E-07 | -0.8422 |
| C1 | SW620 | colon | 1.88E-07 | -0.839 |
| C1 | SU-DHL-6 | hematological | 2.65E-07 | -0.8262 |
| C1 | RAMOS | hematological | 1.59E-07 | -0.8076 |
| C1 | DLD1 | colon | 1.78E-07 | -0.7953 |
| C1 | SKMEL28 | skin | 2.63E-07 | -0.7852 |
| C1 | EJ28 | bladder | 1.72E-07 | -0.7675 |
| C1 | MINO | hematological | 2.47E-07 | -0.7634 |
| C1 | MG63 | bone | 2.39E-07 | -0.7053 |
| C1 | LOVO | colon | 2.62E-07 | -0.6768 |
| C1 | OVCAR4 | ovary | 4.21E-07 | -0.6337 |
| C1 | MDAMB435 | skin | 2.44E-07 | -0.5985 |
| C1 | HT29 | colon | 3.33E-07 | -0.5687 |
| C1 | PANC1 | pancreas | 3.95E-07 | -0.5355 |
| C1 | SKBR3 | breast | 7.89E-07 | -0.5334 |
| C1 | ACHN | kidney | 5.64E-07 | -0.4636 |
| C1 | A431 | skin | 4.26E-07 | -0.4299 |
| C1 | KASUMI-1 | hematological | 4.99E-06 | -0.4286 |
| C1 | HEPG2 | liver | 4.22E-07 | -0.4243 |
| C1 | JIMT1 | breast | 7.04E-07 | -0.3771 |
| C1 | IGROV1 | ovary | 7.14E-07 | -0.3648 |
| C1 | MDAMB436 | breast | - | -0.2949 |
| C1 | CLS439 | bladder | 7.14E-07 | -0.1862 |
| C1 | PLCPRF5 | liver | 5.80E-07 | -0.1766 |
| C1 | DU145 | prostate | 1.39E-06 | 0.0068 |
| C1 | COLO205 | colon | 9.51E-07 | 0.0721 |
| C1 | BXPC3 | pancreas | 2.10E-06 | 0.0855 |
| C1 | NCIH292 | lung | 2.45E-06 | 0.0876 |
| C1 | A673 | muscle | 1.32E-06 | 0.0915 |
| C1 | TE671 | muscle | 9.46E-07 | 0.1128 |
| C1 | PC3 | prostate | 2.14E-06 | 0.1585 |
| C1 | WSU-NHL | hematological | 8.89E-07 | 0.1942 |
| C1 | SKNSH | brain | 4.65E-06 | 0.1985 |
| C1 | NCI-H23 | lung | 1.87E-06 | 0.2372 |
| C1 | ASPC1 | pancreas | 1.37E-06 | 0.2688 |
| C1 | A204 | muscle | 1.71E-06 | 0.3428 |
| C1 | CAKI1 | kidney | 2.13E-06 | 0.3936 |
| C1 | JEG3 | placenta | 1.20E-06 | 0.3991 |
| C1 | CALU6 | lung | 3.86E-06 | 0.5387 |
| C1 | CASKI | endometrial | 2.69E-06 | 0.6131 |
| C1 | OVCAR3 | ovary | 3.22E-06 | 0.6192 |
| C1 | RD | muscle | 5.21E-06 | 0.6516 |
| C1 | MCF7 | breast | 3.27E-06 | 0.6977 |
| C1 | EFO21 | ovary | 8.65E-06 | 0.8272 |
| C1 | 5637 | bladder | 2.73E-06 | 0.8521 |
| C1 | HEK293 | kidney | 3.83E-06 | 0.8724 |
| C1 | 22RV1 | prostate | 2.77E-06 | 0.9083 |
| C1 | SKOV3 | ovary | - | 0.9111 |
| C1 | GRANTA-519 | hematological | 3.95E-06 | 0.9249 |
| C1 | J82 | bladder | 2.79E-06 | 0.9732 |
| C1 | PANC1005 | pancreas | 6.10E-06 | 1.0961 |
| C1 | UO31 | kidney | 4.79E-06 | 1.0962 |
| C1 | CACO2 | colon | 7.00E-06 | 1.1947 |
| C1 | BT20 | breast | - | 1.2169 |
| C1 | SF268 | brain | 9.86E-06 | 1.2301 |
| C1 | SKNAS | brain | - | 1.2744 |
| C1 | NCIH358M | lung | 1.22E-05 | 1.3141 |
| C1 | COLO678 | colon | 5.98E-06 | 1.4093 |
| C1 | SF295 | brain | 7.79E-06 | 1.5228 |
| C1 | MDAMB231 | breast | 1.03E-05 | 1.5422 |
| C1 | IMR90 | lung | - | 1.7451 |
| C1 | U2OS | bone | 1.92E-05 | 1.7557 |
| C1 | SAOS2 | bone | - | 1.7808 |
| C1 | HELA | endometrial | - | 1.8487 |
| C1 | SU-DHL-10 | hematological | - | 2.2091 |
| C1 | U87MG | brain | - | 2.4565 |
| C1 | SNB75 | brain | - | 2.6142 |
| C1 | HS578T | breast | - | - |
| C1 | HS729 | muscle | - | - |
| C1 | PBMC | hematological | - | - |

**Supplementary Table 2.**
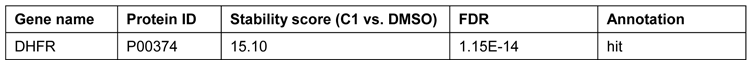
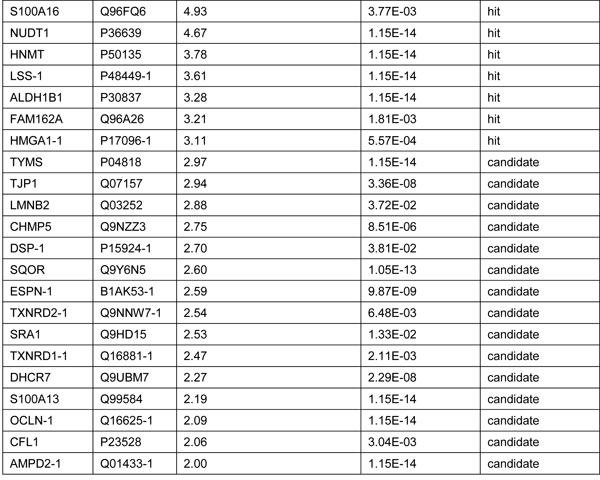
Stabilized hits and candidate hits of compound C1 identified in LS 174T cells, related to Figure 1D. Intact LS 174T cells were treated with 10 μM C1 for 1 hour, followed by thermal proteome profiling. The table of all stabilized hits and candidates includes stability scores and false discovery rates (FDR). Destabilized proteins are excluded from this table but are included as source data.

| Gene name | Protein ID | Stability score (C1 vs. DMSO) | FDR | Annotation |
| --- | --- | --- | --- | --- |
| DHFR | P00374 | 15.10 | 1.15E-14 | hit |
| S100A16 | Q96FQ6 | 4.93 | 3.77E-03 | hit |
| NUDT1 | P36639 | 4.67 | 1.15E-14 | hit |
| HNMT | P50135 | 3.78 | 1.15E-14 | hit |
| LSS-1 | P48449-1 | 3.61 | 1.15E-14 | hit |
| ALDH1B1 | P30837 | 3.28 | 1.15E-14 | hit |
| FAM162A | Q96A26 | 3.21 | 1.81E-03 | hit |
| HMGA1-1 | P17096-1 | 3.11 | 5.57E-04 | hit |
| TYMS | P04818 | 2.97 | 1.15E-14 | candidate |
| TJP1 | Q07157 | 2.94 | 3.36E-08 | candidate |
| LMNB2 | Q03252 | 2.88 | 3.72E-02 | candidate |
| CHMP5 | Q9NZZ3 | 2.75 | 8.51E-06 | candidate |
| DSP-1 | P15924-1 | 2.70 | 3.81E-02 | candidate |
| SQOR | Q9Y6N5 | 2.60 | 1.05E-13 | candidate |
| ESPN-1 | B1AK53-1 | 2.59 | 9.87E-09 | candidate |
| TXNRD2-1 | Q9NNW7-1 | 2.54 | 6.48E-03 | candidate |
| SRA1 | Q9HD15 | 2.53 | 1.33E-02 | candidate |
| TXNRD1-1 | Q16881-1 | 2.47 | 2.11E-03 | candidate |
| DHCR7 | Q9UBM7 | 2.27 | 2.29E-08 | candidate |
| S100A13 | Q99584 | 2.19 | 1.15E-14 | candidate |
| OCLN-1 | Q16625-1 | 2.09 | 1.15E-14 | candidate |
| CFL1 | P23528 | 2.06 | 3.04E-03 | candidate |
| AMPD2-1 | Q01433-1 | 2.00 | 1.15E-14 | candidate |

**Supplementary Table 3.** TMT labels used for thermal proteome profiling.

| TMT label | Condition | Replicate |
| --- | --- | --- |
| 126 | DMSO | 1 |
| 127N | DMSO | 2 |
| 128C | DMSO | 3 |
| 129N | C1 | 1 |
| 130C | C1 | 2 |
| 131 | C1 | 3 |

## Figures and supplementary figures

**Supplementary Figure 1.**
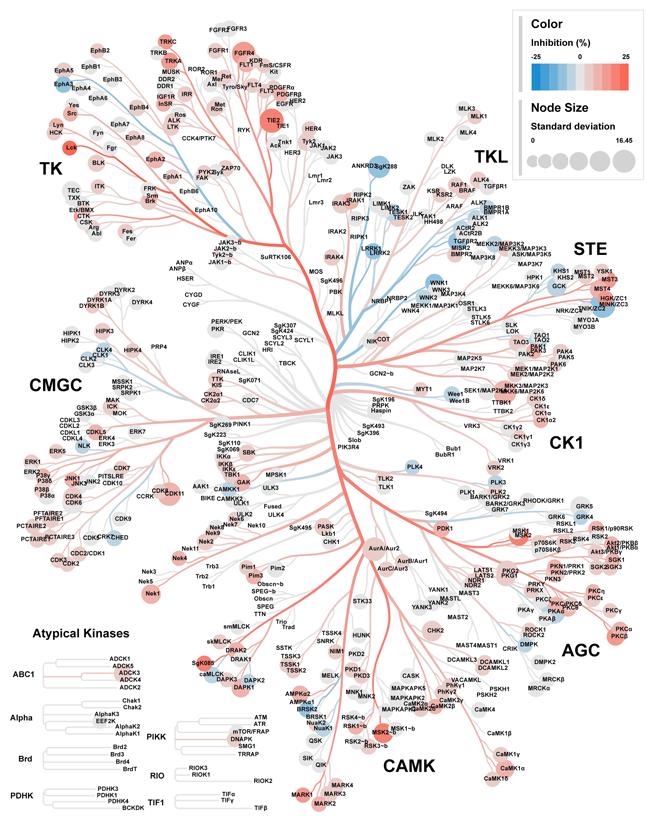
C1 is a poor kinase inhibitor at 100 nM, related to Figure 1. Kinome-wide screening for *in vitro* kinase inhibition by 100 nM C1 treatment, using Adapta, Z’LYTE and LanthaScreen binding assays (see Materials & Methods). Data are mean and s.d., collected at 484 datapoints with n=2 technical replicates. Data were visualized with CORAL (Metz et al., 2018).

**Supplementary Figure 2.**
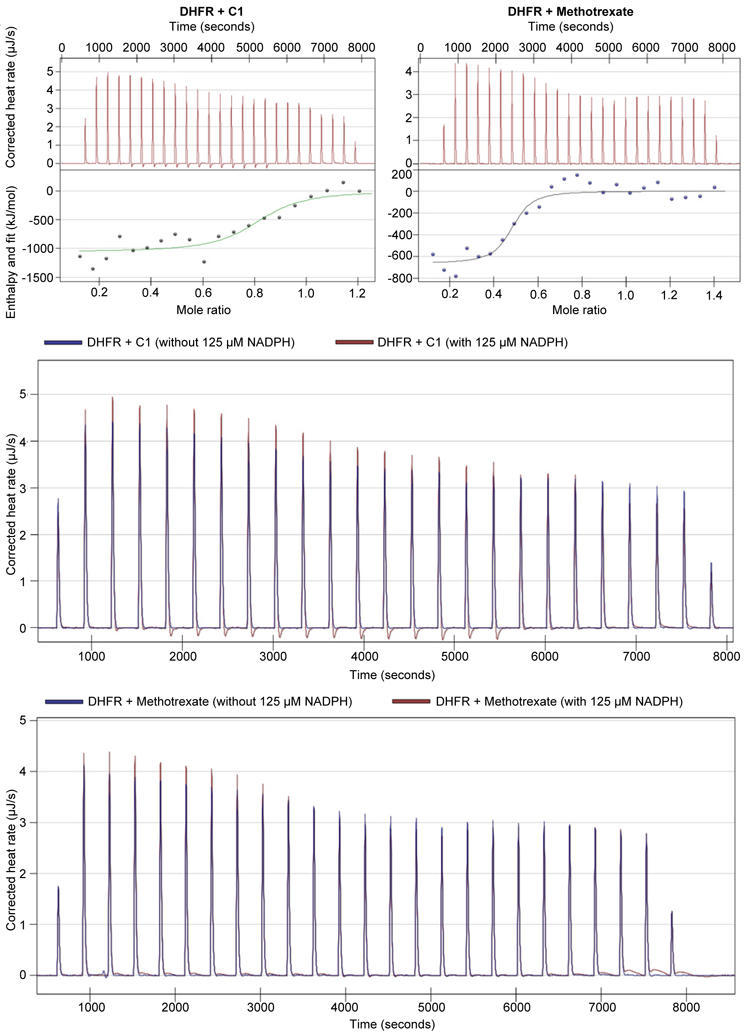
Thermograms of isothermal titration calorimetry experiments, related to Table 1. **(A)** Representative thermograms demonstrating binding of DHFR to C1 and methotrexate. **(B)** Representative thermograms demonstrating that binding of DHFR to C1 and methotrexate is not altered in the presence of 125 μM NADPH.

**Supplementary Figure 3.**
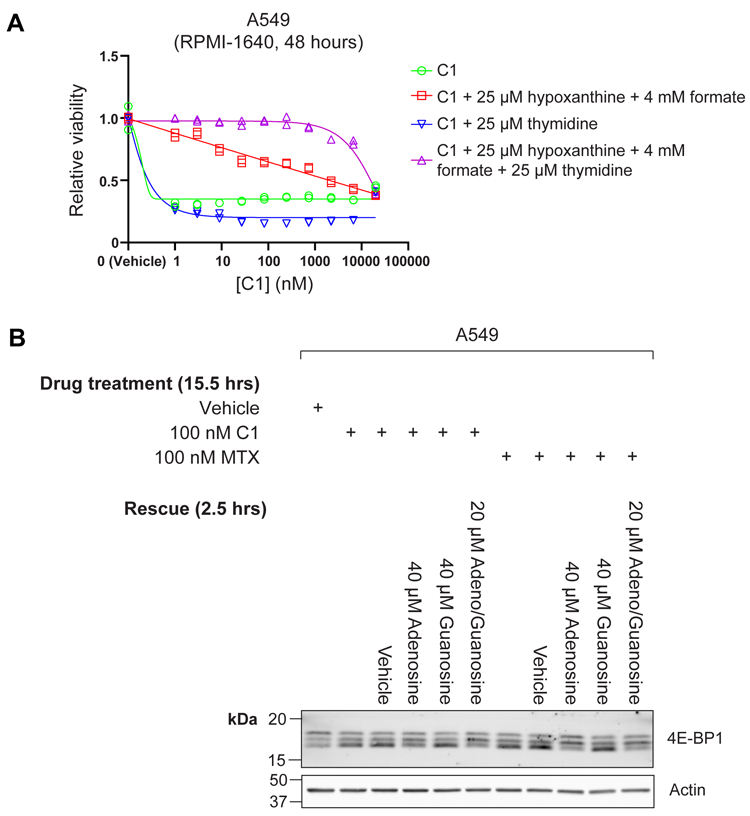
C1 inhibits the folate cycle and modulates purine sensing by mTORC1, similar to methotrexate, related to Figure 2. **(A)** Viability measurement of C1-treated A549 cells after 48 hours treatment, including effects of treatments with indicated products of the folate-mediated one-carbon metabolism. Data was collected in n=2 biological replicates. **(B)** Western-blot analysis of the mTORC1 substrate 4E-BP1 in A549 cells after 15.5 hours 100 nM C1-or 100 nM Methotrexate-treatment, including 2.5 rescue treatments with purines. Rescue treatments consisted of 40 μM adenosine, 40 μM guanosine or a combination of 20 μM adenosine and 20 μM guanosine. Reduced mTORC1 activity leads to reduced 4E-BP1 phosphorylation, which is visualized as a mobility shift to a faster-migrating form.

**Supplementary Figure 4.**
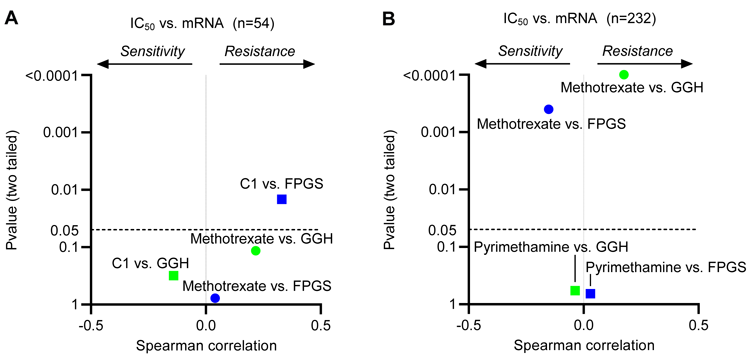
High FPGS expression correlates with resistance to C1, but sensitivity to methotrexate, related toFigure 4. **(A)** Spearman correlation coefficient and p-value between expression of genes involved in folate polyglutamylation and sensitivity to Methotrexate or C1 in a panel of cancer cell lines. Generation of C1 and Methotrexate IC_50_ values is described in Figure 1B. **(B)** Spearman correlation coefficient and p-value between the expression of a selection of genes involved in folate polyglutamylation and sensitivity to Pyrimethamine or Methotrexate for all entries in the Genomics of Drug Sensitivity in Cancer database. Gene expression data were retrieved from the Cancer Cell Line Encyclopedia (Ghandi et al., 2019).

**Supplementary Figure 5.**
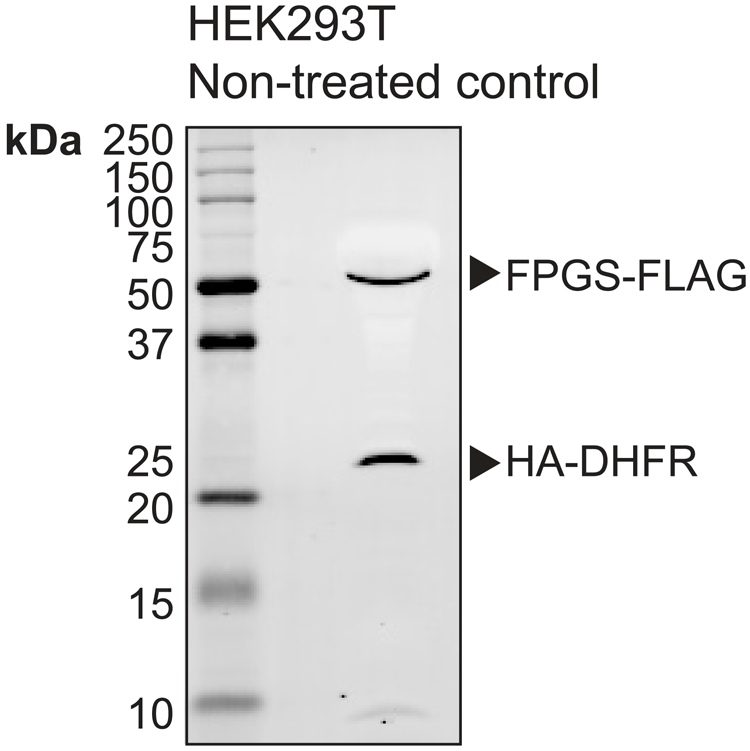
FPGS and DHFR expression in HEK293T cells for intracellular binding studies, related to Figures 1 and 4. Western-blot analysis of overexpressed HA-DHFR and FPGS-FLAG in HEK293T cells used for cellular thermal shift assays in Figures 1 and 4.

**Supplementary Figure 6.**
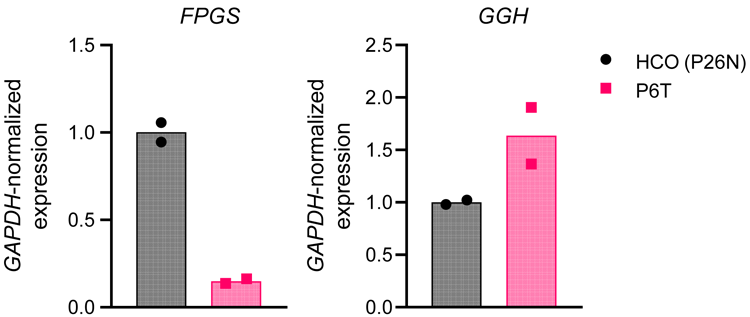
Patient-derived P6T colorectal cancer organoids display altered FPGS and GGH expression compared to healthy colon organoids, related to Figure 5. RT-qPCR analysis of *FPGS* and *GGH* gene expression in healthy colon organoids (HCO) and patient 6-derived (P6T) colorectal cancer organoids. Data were collected in n=2 biological replicates.

**Supplementary Figure 7.**
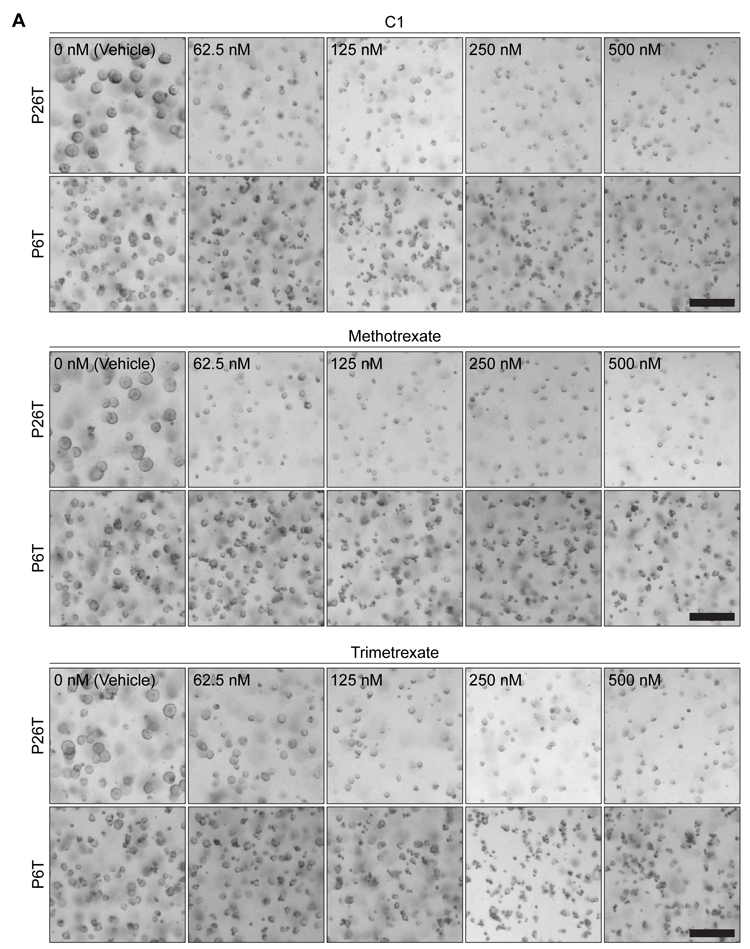
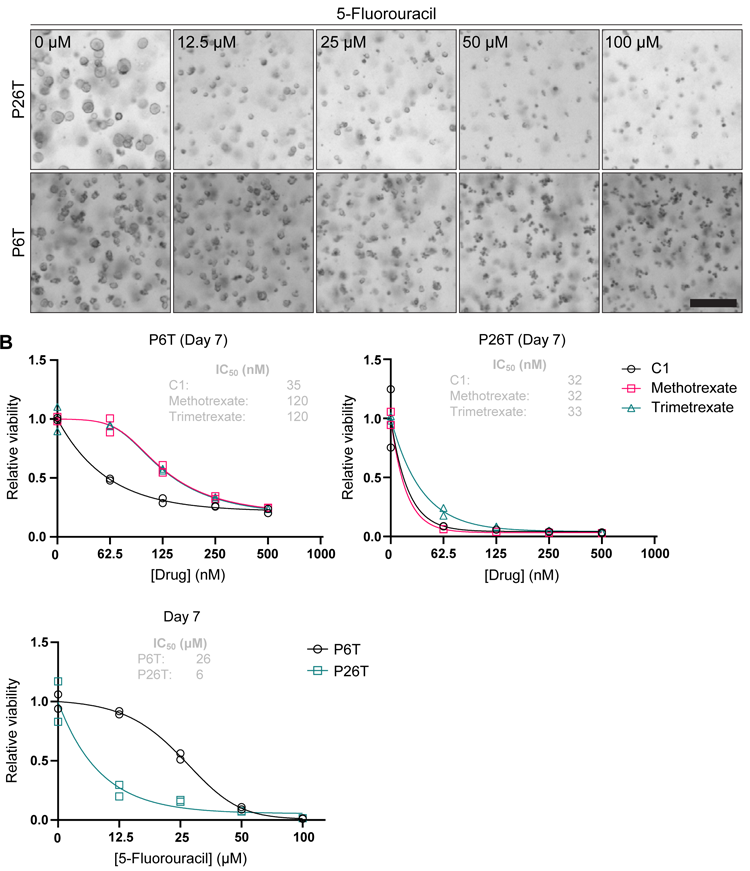
P6T organoids display increased sensitivity to C1, but not trimetrexate, related to Figure 5. **(A)** Brightfield images of P6T and P26T organoids after 7 days of drug treatment. Scalebar represents 250 μm. **(B)** CellTiter-Glo viability analysis of drug-treated P6T and P26T cells after 7 days treatment. Data were collected in n=2 biological replicates.

**Supplementary Figure 8.**
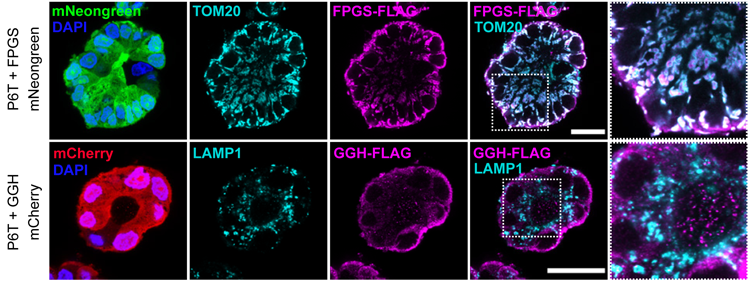
Doxycycline-inducible overexpression of FPGS and GGH in P6T organoids, related to Figure 5. Representative confocal (immuno)fluorescence images of P6T organoids overexpressing FPGS-FLAG IRES-mNeongreen or GGH-FLAG IRES-mCherry in a doxycycline-dependent manner. Organoids were treated with 1 μg/ml doxycycline and stained with anti-FLAG to detect overexpressed proteins, anti-TOM20 to stain mitochondria, anti-LAMP1 to stain lysosomes and DAPI to stain nuclei. Scalebar represents 25 μm.

## References

1. Algul, O., Paulsen, J. L., & Anderson, A. C. (2011). 2, 4-Diamino-5-(2′-arylpropargyl) pyrimidine derivatives as new nonclassical antifolates for human dihydrofolate reductase inhibition. Journal of Molecular Graphics and Modelling, 29(5), 608–613. https://doi.org/10.1016/j.jmgm.2010.11.004

2. Bailey, L. B., Stover, P. J., McNulty, H., Fenech, M. F., Gregory III, J. F., Mills, J. L., … & Raiten, D. J. (2015). Biomarkers of nutrition for development—folate review. The Journal of nutrition, 145(7), 1636S–1680S. https://doi.org/10.3945/jn.114.206599

3. Becher, I., Andrés-Pons, A., Romanov, N., Stein, F., Schramm, M., Baudin, F., Helm, D., Kurzawa, N., Mateus, A., Mackmull, M. T., Typas, A., Müller, C. W., Bork, P., Beck, M., & Savitski, M. M. (2018). Pervasive Protein Thermal Stability Variation during the Cell Cycle. Cell, 173(6), 1495–1507.e18. https://doi.org/10.1016/J.CELL.2018.03.053

4. Becher, I., Werner, T., Doce, C., Zaal, E. A., Tögel, I., Khan, C. A., Rueger, A., Muelbaier, M., Salzer, E., Berkers, C. R., Fitzpatrick, P. F., Bantscheff, M., & Savitski, M. M. (2016). Thermal profiling reveals phenylalanine hydroxylase as an off-target of panobinostat. Nature Chemical Biology 2016 12:11, 12(11), 908–910. https://doi.org/10.1038/nchembio.2185

5. Berman, H. M., Westbrook, J., Feng, Z., Gilliland, G., Bhat, T. N., Weissig, H., Shindyalov, I. N., & Bourne, P. E. (2000). The Protein Data Bank. Nucleic Acids Research, 28(1), 235–242https://doi.org/10.1093/NAR/28.1.235

6. Bhabha, G., Ekiert, D. C., Jennewein, M., Zmasek, C. M., Tuttle, L. M., Kroon, G., Dyson, H. J., Godzik, A., Wilson, I. A., & Wright, P. E. (2013). Divergent evolution of protein conformational dynamics in dihydrofolate reductase. Nature Structural & Molecular Biology, 20(11), 1243–1249. https://doi.org/10.1038/NSMB.2676

7. Brown, J. I., Wang, P., Wong, A. Y., Petrova, B., Persaud, R., Soukhtehzari, S., … & Page, B. D. (2023). Cycloguanil and Analogues Potently Target DHFR in Cancer Cells to Elicit Anti-Cancer Activity. Metabolites, 13(2), 151.

8. Cho, R. C., Cole, P. D., Sohn, K. J., Gaisano, G., Croxford, R., Kamen, B. A., & Kim, Y. I. (2007). Effects of folate and folylpolyglutamyl synthase modulation on chemosensitivity of breast cancer cells. Molecular Cancer Therapeutics, 6(11), 2909–2920. https://doi.org/10.1158/1535-7163.MCT-07-0449

9. Cody, V., Luft, J. R., & Pangborn, W. (2005). Understanding the role of Leu22 variants in methotrexate resistance: comparison of wild-type and Leu22Arg variant mouse and human dihydrofolate reductase ternary crystal complexes with methotrexate and NADPH. Acta Crystallographica. Section D, Biological Crystallography, 61(2), 147–155. https://doi.org/10.1107/S0907444904030422

10. Cuthbertson, C. R., Arabzada, Z., Iii, A. B., Kyani, A., & Neamati, N. (2021). A Review of Small-Molecule Inhibitors of One-Carbon Enzymes: SHMT2 and MTHFD2 in the Spotlight. ACS Pharmacol. Transl. Sci, 2021, 646. https://doi.org/10.1021/acsptsci.0c00223

11. Dominguez, C., Boelens, R., & Bonvin, A. M. J. J. (2003). HADDOCK: A Protein−Protein Docking Approach Based on Biochemical or Biophysical Information. Journal of the American Chemical Society, 125(7), 1731–1737. https://doi.org/10.1021/ja026939x

12. Driehuis, E., Oosterom, N., Heil, S. G., Muller, I. B., Lin, M., Kolders, S., Jansen, G., De Jonge, R., Pieters, R., Clevers, H., & Van Den Heuvel-Eibrink, M. M. (2020). Patient-derived oral mucosa organoids as an in vitro model for methotrexate induced toxicity in pediatric acute lymphoblastic leukemia. PLOS ONE, 15(5), e0231588. https://doi.org/10.1371/JOURNAL.PONE.0231588

13. Ducker, G. S., & Rabinowitz, J. D. (2017). One-Carbon Metabolism in Health and Disease. Cell Metabolism, 25(1), 27–42. https://doi.org/10.1016/j.cmet.2016.08.009

14. Emmanuel, N., Ragunathan, S., Shan, Q., Wang, F., Giannakou, A., Huser, N., Jin, G., Myers, J., Abraham, R. T., & Unsal-Kacmaz, K. (2017). Purine Nucleotide Availability Regulates mTORC1 Activity through the Rheb GTPase. Cell Reports, 19(13), 2665–2680. https://doi.org/10.1016/j.celrep.2017.05.043

15. Fabre, I., Fabre, G., & Goldman, I. D. (1984). Polyglutamylation, an Important Element in Methotrexate Cytotoxicity and Selectivity in Tumor versus Murine Granulocytic Progenitor Cells in Vitro. Cancer Research, 44(8), 3190–3195.

16. Fernández-Recio, J., Totrov, M., & Abagyan, R. (2004). Identification of Protein–Protein Interaction Sites from Docking Energy Landscapes. Journal of Molecular Biology, 335(3), 843–865. https://doi.org/10.1016/j.jmb.2003.10.069

17. Fotoohi, A. K., Assaraf, Y. G., Moshfegh, A., Hashemi, J., Jansen, G., Peters, G. J., Larsson, C., & Albertioni, F. (2009). Gene expression profiling of leukemia T-cells resistant to methotrexate and 7-hydroxymethotrexate reveals alterations that preserve intracellular levels of folate and nucleotide biosynthesis. Biochemical Pharmacology, 77(8), 1410–1417https://doi.org/10.1016/J.BCP.2008.12.026

18. Fujii, M., Matano, M., Nanki, K., & Sato, T. (2015). Efficient genetic engineering of human intestinal organoids using electroporation. Nature Protocols 10(10), 1474–1485. https://doi.org/10.1038/nprot.2015.088

19. Gangjee, A., Jain, H. D., & Kurup, S. (2007). Recent Advances in Classical and Non-Classical Antifolates as Antitumor and Antiopportunistic Infection Agents: Part I. Anti-Cancer Agents in Medicinal Chemistry, 7(5), 524–542. https://doi.org/10.2174/187152007781668724

20. Gangjee, A., Jain, H., & Kurup, S. (2008). Recent Advances in Classical and Non-Classical Antifolates as Antitumor and Antiopportunistic Infection Agents: Part II. Anti-Cancer Agents in Medicinal Chemistry, 8(2), 205–231. https://doi.org/10.2174/187152008783497064

21. Ghandi, M., Huang, F. W., Jané-Valbuena, J., Kryukov, G. V., Lo, C. C., McDonald, E. R., Barretina, J., Gelfand, E. T., Bielski, C. M., Li, H., Hu, K., Andreev-Drakhlin, A. Y., Kim, J., Hess, J. M., Haas, B. J., Aguet, F., Weir, B. A., Rothberg, M. V., Paolella, B. R., … Sellers, W. R. (2019). Next-generation characterization of the Cancer Cell Line Encyclopedia. Nature, 569(7757), 503–508. https://doi.org/10.1038/S41586-019-1186-3

22. Hoxhaj, G., Hughes-Hallett, J., Timson, R. C., Ilagan, E., Yuan, M., Asara, J. M., Ben-Sahra, I., & Manning, B. D. (2017). The mTORC1 Signaling Network Senses Changes in Cellular Purine Nucleotide Levels. Cell Reports, 21(5), 1331–1346. https://doi.org/10.1016/j.celrep.2017.10.029

23. Huber, K. V. M., Olek, K. M., Müller, A. C., Tan, C. S. H., Bennett, K. L., Colinge, J., & Superti-Furga, G. (2015). Proteome-wide drug and metabolite interaction mapping by thermal-stability profiling. Nature Methods, 12(11), 1055–1057. https://doi.org/10.1038/nmeth.3590

24. Huber, W., Von Heydebreck, A., Sültmann, H., Poustka, A., & Vingron, M. (2002). Variance stabilization applied to microarray data calibration and to the quantification of differential expression. Bioinformatics, 18 (SUPPL_1), S96–S104. https://doi.org/10.1093/BIOINFORMATICS/18.SUPPL_1.S96

25. Jorgensen, W. L., & Tirado-Rives, J. (1988). The OPLS [optimized potentials for liquid simulations] potential functions for proteins, energy minimizations for crystals of cyclic peptides and crambin. Journal of the American Chemical Society, 110(6), 1657–1666. https://doi.org/10.1021/ja00214a001

26. Kim, S. E., Cole, P. D., Cho, R. C., Ly, A., Ishiguro, L., Sohn, K. J., Croxford, R., Kamen, B. A., & Kim, Y. I. (2013). γ-Glutamyl hydrolase modulation and folate influence chemosensitivity of cancer cells to 5-fluorouracil and methotrexate. British Journal of Cancer, 109(8), 2175–2188. https://doi.org/10.1038/bjc.2013.579

27. Koukos, P. I., Réau, M., & Bonvin, A. M. J. J. (2021). Shape-Restrained Modeling of Protein-Small-Molecule Complexes with High Ambiguity Driven DOCKing. Journal of Chemical Information and Modeling, 61(9), 4807–4818. https://doi.org/10.1021/ACS.JCIM.1C00796/SUPPL_FILE/CI1C00796_SI_001.PDF

28. Lawrence, S. A., Titus, S. A., Ferguson, J., Heineman, A. L., Taylor, S. M., & Moran, R. G. (2014). Mammalian mitochondrial and cytosolic folylpolyglutamate synthetase maintain the subcellular compartmentalization of folates. Journal of Biological Chemistry. https://doi.org/10.1074/jbc.M114.593244

29. Li, B., Brady, S. W., Ma, X., Shen, S., Zhang, Y., Li, Y., Szlachta, K., Dong, L., Liu, Y., Yang, F., Wang, N., Flasch, D. A., Myers, M. A., Mulder, H. L., Ding, L., Liu, Y., Tian, L., Hagiwara, K., Xu, K., … Zhang, J. (2020). Therapy-induced mutations drive the genomic landscape of relapsed acute lymphoblastic leukemia. Blood, 135(1), 41–55. https://doi.org/10.1182/BLOOD.2019002220

30. Liani, E., Rothem, L., Bunni, M. A., Smith, C. A., Jansen, G., & Assaraf, Y. G. (2003). Loss of folylpoly-γ-glutamate synthetase activity is a dominant mechanism of resistance to polyglutamylation-dependent novel antifolates in multiple human leukemia sublines. International Journal of Cancer, 103(5), 587–599. https://doi.org/10.1002/ijc.10829

31. Longley, D. B., Harkin, D. P., & Johnston, P. G. (2003). 5-Fluorouracil: mechanisms of action and clinical strategies. Nature Reviews Cancer, 3(5), 330–338. https://doi.org/10.1038/nrc1074

32. Mateus, A., Hevler, J., Bobonis, J., Kurzawa, N., Shah, M., Mitosch, K., Goemans, C. V, Helm, D., Stein, F., Typas, A., & Savitski, M. M. (2020). The functional proteome landscape of Escherichia coli. Nature, 588, 473–478. https://doi.org/10.1038/s41586-020-3002-5

33. Mateus, A., Kurzawa, N., Becher, I., Sridharan, S., Helm, D., Stein, F., Typas, A., & Savitski, M. M. (2020). Thermal proteome profiling for interrogating protein interactions. Molecular Systems Biology, 16(3), e9232. https://doi.org/10.15252/MSB.20199232

34. Metz, K. S., Deoudes, E. M., Berginski, M. E., Jimenez-Ruiz, I., Aksoy, B. A., Hammerbacher, J., Gomez, S. M., & Phanstiel, D. H. (2018). Coral: Clear and Customizable Visualization of Human Kinome Data. Cell Systems, 7(3), 347–350.e1. https://doi.org/10.1016/J.CELS.2018.07.001

35. Mi, H., Muruganujan, A., Huang, X., Ebert, D., Mills, C., Guo, X., & Thomas, P. D. (2019). Protocol Update for large-scale genome and gene function analysis with the PANTHER classification system (v.14.0). Nature Protocols, 14(3), 703–721. https://doi.org/10.1038/S41596-019-0128-8

36. Molina, D. M., Jafari, R., Ignatushchenko, M., Seki, T., Larsson, E. A., Dan, C., Sreekumar, L., Cao, Y., & Nordlund, P. (2013). Monitoring drug target engagement in cells and tissues using the cellular thermal shift assay. Science, 341(6141), 84–87. https://doi.org/10.1126/SCIENCE.1233606

37. Nguyen, K., Devidas, M., Cheng, S. C., La, M., Raetz, E. A., Carroll, W. L., Winick, N. J., Hunger, S. P., Gaynon, P. S., & Loh, M. L. (2008). Factors influencing survival after relapse from acute lymphoblastic leukemia: a Children’s Oncology Group study. Leukemia, 22(12), 2142–2150. https://doi.org/10.1038/LEU.2008.251

38. Nilsson, R., Jain, M., Madhusudhan, N., Sheppard, N. G., Strittmatter, L., Kampf, C., … & Mootha, V. K. (2014). Metabolic enzyme expression highlights a key role for MTHFD2 and the mitochondrial folate pathway in cancer. Nature communications, 5(1), 3128. https://doi.org/10.1038/ncomms4128

39. Omerzu, M., Fenderico, N., De Barbanson, B., Sprangers, J., De Ridder, J., & Maurice, M. M. (2019). Three-dimensional analysis of single molecule FISH in human colon organoids. Biology Open, 8(8). https://doi.org/10.1242/BIO.042812

40. Osborne, C. B., Lowe, K. E., & Shane, B. (1993). Regulation of folate and one-carbon metabolism in mammalian cells. I. Folate metabolism in Chinese hamster ovary cells expressing Escherichia coli or human folylpoly-gamma-glutamate synthetase activity. Journal of Biological Chemistry, 268(29), 21657–21664. https://doi.org/10.1016/S0021-9258(20)80592-4

41. Perez-Riverol, Y., Csordas, A., Bai, J., Bernal-Llinares, M., Hewapathirana, S., Kundu, D. J., … & Vizcaíno, J. A. (2019). The PRIDE database and related tools and resources in 2019: improving support for quantification data. Nucleic acids research, 47(D1), D442–D450. https://doi.org/10.1093/nar/gky1106

42. Piper, J. R., McCaleb, G. S., Montgomery, J. A., Kisliuk, R. L., Gaumont, Y., & Sirotnak, F. M. (1985). Syntheses and antifolate activity of 5-methyl-5-deaza analogs of aminopterin, methotrexate, folic acid, and N10-methylfolic acid. Journal of Medicinal Chemistry, 29(6), 1080–1087. https://doi.org/10.1021/JM00156A029

43. Rappsilber, J., Mann, M., & Ishihama, Y. (2007). Protocol for micro-purification, enrichment, pre-fractionation and storage of peptides for proteomics using StageTips. Nature Protocols, 2(8), 1896–1906. https://doi.org/10.1038/NPROT.2007.261

44. Riniker, S., & Landrum, G. A. (2015). Better Informed Distance Geometry: Using What We Know To Improve Conformation Generation. Journal of Chemical Information and Modeling, 55(12), 2562–2574. https://doi.org/10.1021/acs.jcim.5b00654

45. Ritchie, M. E., Phipson, B., Wu, D., Hu, Y., Law, C. W., Shi, W., & Smyth, G. K. (2015). Limma powers differential expression analyses for RNA-sequencing and microarray studies. Nucleic Acids Research, 43(7), e47. https://doi.org/10.1093/NAR/GKV007

46. Rots, M. G., Pieters, R., Peters, G. J., Noordhuis, P., Van Zantwijk, C. H., Kaspers, G. J. L., Hählen, K., Creutzig, U., Veerman, A. J. P., & Jansen, G. (1999). Role of Folylpolyglutamate Synthetase and Folylpolyglutamate Hydrolase in Methotrexate Accumulation and Polyglutamylation in Childhood Leukemia. Blood, 93(5), 1677–1683. https://doi.org/10.1182/BLOOD.V93.5.1677

47. Savitski, M. M., Reinhard, F. B. M., Franken, H., Werner, T., Savitski, M. F., Eberhard, D., Molina, D. M., Jafari, R., Dovega, R. B., Klaeger, S., Kuster, B., Nordlund, P., Bantscheff, M., & Drewes, G. (2014). Tracking cancer drugs in living cells by thermal profiling of the proteome. Science, 346(6205), 1255784. https://doi.org/10.1126/SCIENCE.1255784

48. Schüttelkopf, A. W., & Van Aalten, D. M. (2004). PRODRG: a tool for high-throughput crystallography of protein–ligand complexes. Acta Crystallographica Section D: Biological Crystallography, 60(8), 1355–1363. https://doi.org/10.1107/s0907444904011679

49. Sohn, K. J., Smirnakis, F., Moskovitz, D. N., Novakovic, P., Yates, Z., Lucock, M., Croxford, R., & Kim, Y. I. (2004). Effects of folylpolyglutamate synthetase modulation on chemosensitivity of colon cancer cells to 5-fluorouracil and methotrexate. Gut, 53(12), 1825–1831. https://doi.org/10.1136/gut.2004.042713

50. Stark, M., Wichman, C., Avivi, I., & Assaraf, Y. G. (2009). Aberrant splicing of folylpolyglutamate synthetase as a novel mechanism of antifolate resistance in leukemia. Blood, 113(18), 4362–4369. https://doi.org/10.1182/blood-2008-08-173799

51. Stine, Z. E., Schug, Z. T., Salvino, J. M., & Dang, C. V. (2022). Targeting cancer metabolism in the era of precision oncology. Nature Reviews Drug Discovery, 21(2), 141–162. https://doi.org/10.1038/S41573-021-00339-6

52. Takami, M., Kuniyoshi, Y., Oomukai, T., Ishida, T., & Yamano, Y. (1995). Severe Complications After High-Dose Methotrexate Treatment. Acta Oncologica, 34(5), 611–612. https://doi.org/10.3109/02841869509094036

53. Tversky, A. (1977). Features of similarity. Psychological Review, 84(4), 327–352. https://doi.org/10.1037/0033-295X.84.4.327

54. Van de Wetering, M., Francies, H. E., Francis, J. M., Bounova, G., Iorio, F., Pronk, A., Van Houdt, W., Van Gorp, J., Taylor-Weiner, A., Kester, L., McLaren-Douglas, A., Blokker, J., Jaksani, S., Bartfeld, S., Volckman, R., Van Sluis, P., Li, V. S. W., Seepo, S., Sekhar Pedamallu, C., … Clevers, H. (2015). Prospective derivation of a living organoid biobank of colorectal cancer patients. Cell, 161(4), 933–945. https://doi.org/10.1016/J.CELL.2015.03.053

55. Van der Velden, L. M., Maas, P., van Amersfoort, M., Timmermans-Sprang, E. P. M., Mensinga, A., van der Vaart, E., Malergue, F., Viëtor, H., Derksen, P. W. B., Klumperman, J., van Agthoven, A., Egan, D. A., Mol, J. A., & Strous, G. J. (2022). Small molecules to regulate the GH/IGF1 axis by inhibiting the growth hormone receptor synthesis. Frontiers in Endocrinology, 13, 1631. https://doi.org/10.3389/FENDO.2022.926210/BIBTEX

56. Van Zundert, G. C. P., Rodrigues, J. P. G. L. M., Trellet, M., Schmitz, C., Kastritis, P. L., Karaca, E., Melquiond, A. S. J., van Dijk, M., de Vries, S. J., & Bonvin, A. M. J. J. (2016). The HADDOCK2.2 Web Server: User-Friendly Integrative Modeling of Biomolecular Complexes. Journal of Molecular Biology, 428(4), 720–725. https://doi.org/10.1016/j.jmb.2015.09.014

57. Wang, S., Witek, J., Landrum, G. A., & Riniker, S. (2020). Improving Conformer Generation for Small Rings and Macrocycles Based on Distance Geometry and Experimental Torsional-Angle Preferences. Journal of Chemical Information and Modeling, 60(4), 2044–2058. https://doi.org/10.1021/acs.jcim.0c00025

58. Weininger, D. (1988). SMILES, a chemical language and information system. 1. Introduction to methodology and encoding rules. Journal of Chemical Information and Modeling, 28(1), 31–36. https://doi.org/10.1021/ci00057a005

59. Weininger, D., Weininger, A., & Weininger, J. L. (1989). SMILES. 2. Algorithm for generation of unique SMILES notation. Journal of Chemical Information and Computer Sciences, 29(2), 97–101. https://doi.org/10.1021/ci00062a008

60. Wilson, P. M., Danenberg, P. V., Johnston, P. G., Lenz, H.-J., & Ladner, R. D. (2014). Standing the test of time: targeting thymidylate biosynthesis in cancer therapy. Nature Reviews Clinical Oncology, 11(5), 282–298. https://doi.org/10.1038/nrclinonc.2014.51

61. Wojtuszkiewicz, A., Raz, S., Stark, M., Assaraf, Y. G., Jansen, G., Peters, G. J., Sonneveld, E., Kaspers, G. J. L., & Cloos, J. (2016). Folylpolyglutamate synthetase splicing alterations in acute lymphoblastic leukemia are provoked by methotrexate and other chemotherapeutics and mediate chemoresistance. International Journal of Cancer, 138(7), 1645–1656. https://doi.org/10.1002/IJC.29919

62. Yang, M., & Vousden, K. H. (2016). Serine and one-carbon metabolism in cancer. Nature Reviews Cancer, 16(10), 650–662. https://doi.org/10.1038/nrc.2016.81

63. Yu, S. L., Zhang, H., Ho, B. C., Yu, C. H., Chang, C. C., Hsu, Y. C., Ni, Y. L., Lin, K. H., Jou, S. T., Lu, M. Y., Chen, S. H., Wu, K. H., Wang, S. C., Chang, H. H., Pui, C. H., Yang, J. J., Zhang, J., Lin, D. T., Lin, S. W., … Yang, Y. L. (2020). FPGS relapse-specific mutations in relapsed childhood acute lymphoblastic leukemia. Scientific Reports, 10(1), 1–8. https://doi.org/10.1038/S41598-020-69059-Y

64. Zarou, M. M., Vazquez, A., & Vignir Helgason, G. (2021). Folate metabolism: a re-emerging therapeutic target in haematological cancers. Leukemia, 35(6), 1539–1551. https://doi.org/10.1038/s41375-021-01189-2

65. Zhao, R., Diop-Bove, N., Visentin, M., & Goldman, I. D. (2011). Mechanisms of Membrane Transport of Folates into Cells and Across Epithelia. Annual review of nutrition, 31, 177–201. https://doi.org/10.1146/ANNUREV-NUTR-072610-145133

66. Zhao, R., & Goldman, I. D. (2003). Resistance to antifolates. Oncogene, 22(47), 7431–7457. https://doi.org/10.1038/sj.onc.1206946

67. Zhao, R., Min, S. H., Wang, Y., Campanella, E., Low, P. S., & Goldman, I. D. (2009). A Role for the Proton-coupled Folate Transporter (PCFT-SLC46A1) in Folate Receptor-mediated Endocytosis. Journal of Biological Chemistry, 284(7), 4267–4274. https://doi.org/10.1074/JBC.M807665200

68. Zheng, Y., & Cantley, L. C. (2019). Toward a better understanding of folate metabolism in health and disease. Journal of Experimental Medicine, 216(2), 253–266. https://doi.org/10.1084/jem.20181965

69. Zheng, Y., Lin, T. Y., Lee, G., Paddock, M. N., Momb, J., Cheng, Z., Li, Q., Fei, D. L., Stein, B. D., Ramsamooj, S., Zhang, G., Blenis, J., & Cantley, L. C. (2018). Mitochondrial One-Carbon Pathway Supports Cytosolic Folate Integrity in Cancer Cells. Cell, 175(6), 1546–1560. https://doi.org/10.1016/j.cell.2018.09.041

